# Resource acquisition is more sensitive than carbon storage in soil microorganisms under climate extremes

**DOI:** 10.64898/2026.08.03.742454

**Authors:** Itzel Lopez-Montoya, Qingdong Zhu, Ludovico Formenti, Nicolo Tartini, Anita C. Risch, Irene Cordero, Nicholas O.E. Ofiti, Madhav P. Thakur

## Abstract

1. Drought and warming can disrupt soil microbial processes and ecosystem functioning. Although soil microorganisms can exhibit physiological adjustments to drought, it remains unclear how they allocate resources between extracellular resource acquisition, potential oxidative metabolism, and carbon storage during drought and recovery, particularly under constant warming and/or heat waves.
2. Here, we tested the effects of drought on microbial resource allocation strategies across warming regimes during the resistance and recovery phases. We performed a full-factorial outdoor mesocosm experiment combining drought with constant warming and periodic heat waves, applied individually and in combination. We measured the potential activities of extracellular enzymes as proxy for the acquisition of microbial resources, the activity of dehydrogenase as a proxy for the potential active oxidative metabolism, and microbial glycogen pools as a proxy for carbon storage. We also quantified drought legacy effects by measuring microbial functioning before the new drought treatments, capturing the influence of the drought imposed in the previous year.
3. During the resistance phase, dehydrogenase activity and glycogen pools remained stable, despite reduced extracellular enzyme production, while enzyme allocation shifted towards oxidative enzymes associated with acquisition of recalcitrant C in warming regimes. One month after rewetting, all microbial proxies no longer differed from the control soil moisture conditions. Drought legacy effects were observed in extracellular enzymes, dehydrogenase activity, and glycogen pools, with glycogen exhibiting the strongest legacy effect.
4. We conclude that the asymmetrical responses of extracellular resource acquisition and internal C storage to drought and warming may function as strategies for microbial survival in increasingly variable climates.

## 1. Introduction

Compound climate extremes such as drought, chronic warming, and periodic heat waves are increasing in frequency and intensity (AghaKouchak et al., 2020; IPCC, 2023). These events pose a significant threat to soil microorganisms and functions (Bardgett & Caruso, 2020; Jansson & Hofmockel, 2020). Drought and warming impose contrasting constraints on microbial composition, survival, and growth: drought limits pore diffusion and resource availability, while warming increases microbial metabolic rates and resource demands, particularly when water and substrate availability are not limiting (Schimel et al., 2007; Schimel, 2018; Knight et al., 2024). In response to these environmental constraints, microorganisms can reallocate energy between functions associated with resource acquisition, active oxidative metabolism, and internal C storage, thus balancing growth and survival under stress (Malik et al., 2020; Manzoni et al., 2021; Butler et al., 2023; Bouskill et al., 2024). Furthermore, drought can generate legacy effects that alter microbial responses to subsequent climate events, through changes in community composition or functional potential (Canarini et al., 2021; Müller & Bahn, 2022). Thus, despite restored soil diffusion and substrate availability after rewetting, microbial recovery trajectories may vary widely depending on the intensity and frequency of the stressor, and by substrate availability, ranging from incomplete recovery (under-recovery), full recovery, or even overcompensation (over-recovery) (Bardgett & Caruso, 2020; Thakur et al., 2022). However, how compound extremes shape microbial activities, allocation strategies, and their recovery trajectories remains poorly understood, particularly under the combined effects of drought, warming, and heat waves (Bardgett & Caruso, 2020; Thakur et al., 2022; Allison, 2023).

Soil microbial functional activities and resource allocation strategies under climate extremes can be inferred from extracellular enzyme activities as proxies for resource acquisition, dehydrogenase activity (DEH) as an indicator of active oxidative metabolism linked to microbial respiration and ATP production, and glycogen concentration (GLY) as one of the measures for microbial internal carbon (C) storage (Sinsabaugh, 1994; Camiña et al., 1998; Mason-Jones et al., 2022). Specifically, extracellular enzymes include hydrolases and oxidases associated with labile and recalcitrant C degradation, respectively, as well as enzymes involved in the acquisition of nitrogen (N) and phosphorus (P). The capacity to shift among these allocation strategies is constrained by community diversity, soil properties, resource availability, and the intensity, duration, and frequency of climatic stressors (Burns et al., 2013; Wallenstein & Hall, 2012; Domeignoz-Horta et al., 2023).

Drought reduces microbial resource acquisition (Xiao et al., 2018; Cordero et al., 2023), increases investment in internal storage and maintenance processes (Canarini et al., 2024). Additionally, the legacy effects of drought can persist into the following year, reducing resource acquiring enzymes and shifting the microbial allocation away from N acquisition toward oxidative enzyme production (Canarini et al., 2021; Oram et al., 2025). Furthermore, studies have shown potential extracellular enzyme activities after intense drought, while fully recovery has been observed after milder droughts (Cordero et al., 2023), or over-recovery after three years of drying-rewetting cycles (Hammerl et al., 2019). These drying-rewetting cycles can increase microbial activity through resource accumulation and labile C release through physical disruption, although repeated cycles may deplete substrates and alter ecosystem functioning (Fanin et al., 2022; Domeignoz-Horta et al., 2023). On the contrary, warming generally increases microbial growth, nutrient demand, and extracellular enzyme activity, while increasing investment in oxidases as microbes shift from labile to more complex C substrates following resource depletion (Sinsabaugh, 2010; Hassan et al., 2013; Fanin et al., 2022; Domeignoz-Horta et al., 2023). Additionally, the warming rate can redirect the allocation of microbial enzymes: gradual warming increased microbial carbon use efficiency (CUE) and microbial biomass carbon (MBC), together with higher enzyme activities that acquire N and P, while step warming reduced CUE and MBC while increasing C-acquiring enzyme activity, suggesting microbial acclimation under gradual warming, and greater metabolic stress or microbial community shifts under step warming (Sihi et al., 2019). Studies that examine drought x warming interactions have reported changes in microbial resource allocation, including reduced N acquisition and greater investment in recalcitrant C acquisition (Zhu et al., 2021), although others found no interactive effects on extracellular enzyme activities (Steinweg et al., 2013). These drought x warming interactions on extracellular enzyme activities are highly variable, depending on soil moisture, substrate, nutrient availability, and plant-mediated responses (Zuccarini et al., 2023). Therefore, the evidence for microbial functional activities and allocation strategies to compound drought and warming remains inconsistent, particularly from field studies evaluating the effects of heat waves, during the legacy, resistance, and recovery phases (Fanin et al., 2022; Müller & Bahn, 2022; Thakur et al., 2022; Zuccarini et al., 2023).

In an outdoor mesocosm experiment, we determined the effects of drought on soil microbial functional activities under constant warming, heat waves, and their combination at three temporal stages, including the effects of drought of the summer drought of the previous year in spring before the onset of the summer drought (drought legacy), immediately after drought (resistance) and one month after rewetting (recovery) (Fig. 1). We quantify microbial activities by measuring extracellular enzymes, dehydrogenase activity, and glycogen pools as indicators of potential resource acquisition, potential active oxidative metabolism, and internal C storage, respectively. We hypothesize that 1) the legacy of drought will reduce resource acquisition and during the resistance phase it will reduce resource acquiring enzymes, particularly labile C acquisition, while increasing microbial allocation to internal C storage and maintaining potential oxidative metabolism (Canarini et al., 2021, 2024). These functional shifts are expected to persist after rewetting, resulting in under-recovery of resource acquiring enzymes (Müller & Bahn, 2022; Cordero et al., 2023); 2) warming will amplify drought constraints by increasing potential enzyme allocation towards recalcitrant C acquisition, with heat waves imposing greater metabolic constraints during the resistance phase, followed by over-recovery under these two warming regimes (Sihi et al., 2019; Zhu et al., 2021), 3) drought combined with constant warming and heat waves will impose strongest functional constraints, resulting in greatest reductions in resource acquiring enzymes, while maintaining potential oxidative metabolism during the resistance phase, and continuing to cause under-recovery (Zuccarini et al., 2023).

**Figure 1.**
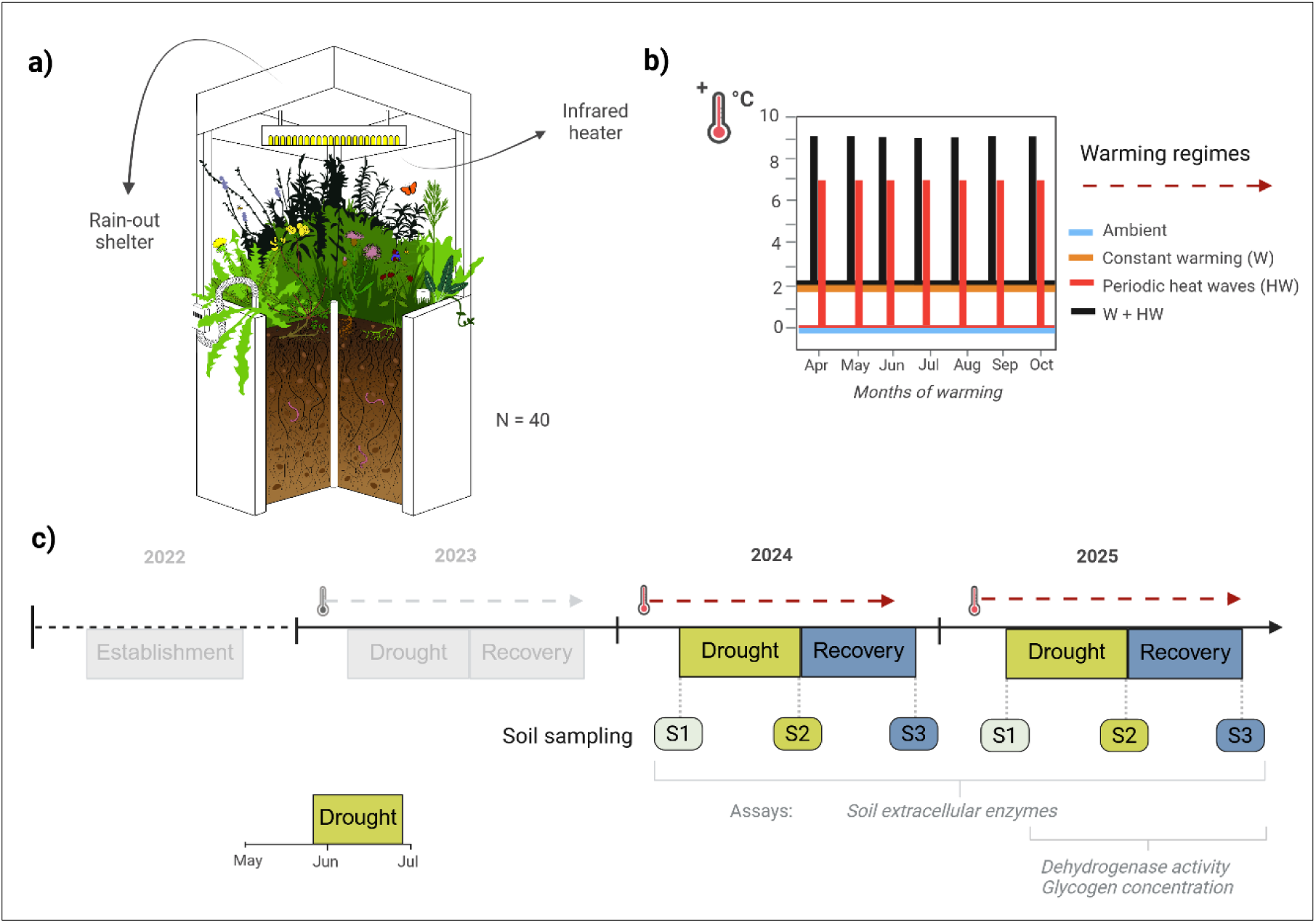
Experimental design of the multifactorial mesocosm experiment. a) Experimental units consisting of 40 mesocosms, a rain-out shelter, and an infrared heater to simulate drought and warming, respectively. b) Warming regimes: 1) ‘Ambient’, 2) ‘Constant warming’ (W), 3) ‘Heat Waves’ (HW), and 4) ‘W + HW’; c) Experimental timeline and sampling scheme. The experiment was established in 2022, treatments began in 2023, and soil sampling was conducted in 2024 and 2025. In each year, soils were sampled repeatedly at three time points: ‘drought legacy phase’ (S1), ‘resistance phase’ (S2), and ‘recovery phase’ (S3). Drought and warming treatments were applied annually during the growing season.

## 2. Materials and Methods

Replication statement.

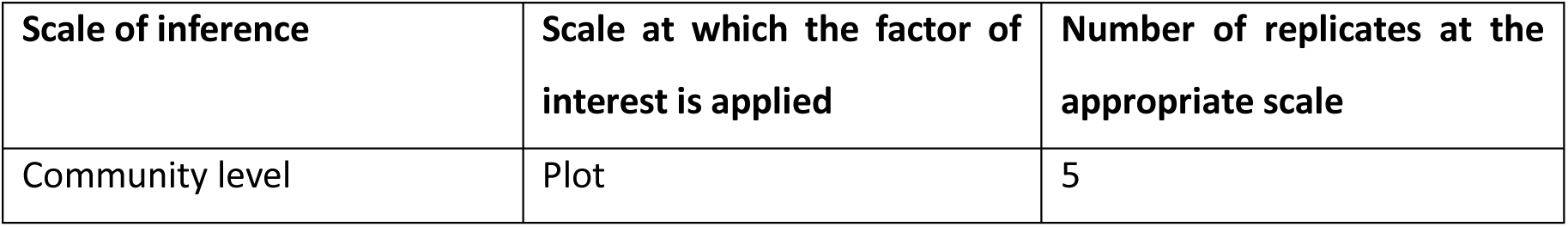

| Scale of inference | Scale at which the factor of interest is applied | Number of replicates at the appropriate scale |
| --- | --- | --- |
| Community level | Plot | 5 |

### 2.1 Experimental site, mesocosm design, and soil sampling

The study was carried out using an outdoor mesocosm experiment located at the Hasli Ethological Station, Bern, Switzerland (46 ° 58′01.362′ N, 7°23′50.243″E). Each mesocosm measured 1 x 1 x 0.45 m (L x W x H). The mesocosms were filled with a custom-formulated commercial soil mixture designed specifically to replicate the physicochemical properties of mineral soil from a nearby managed meadow. Each mesocosm was then topped with a 5-cm layer of homogenised and sieved organic matter-rich topsoil collected from the surrounding grassland. This material, which had a pH of 7.21, served as a live soil inoculum. The baseline soil was classified as silt loam, comprising 6.7% clay (<2.0 μm), 50.3% silt (2–63 μm), and 43.0% sand (63–2000 μm). Its initial chemical properties were as follows: a pH of 7.09, ∼15.71% organic matter (determined by loss of ignition at 550°C), ∼0.49% total N, 75.89 mg PO₄ kg⁻¹ dry soil, 7.90 NO₃^-^ mg kg^-1^ dry soil, 1.66 NH ^+^ mg kg^-1^ dry soil. This mesocosm design provided a balance between experimental control and ecological realism, allowing treatment effects to be assessed under standardized conditions while retaining key characteristics of the surrounding meadow ecosystem (Stewart et al., 2013). All mesocosms contained the same plant community (representative of extensively managed central European meadows), consisting of eight native species *(Lotus corniculatus, Trifolium pratense, Centaurea jacea, Taraxacum officinale, Bromus erectus, Holcus lanatus, Salvia pratensis, Prunella vulgaris*) and one invasive species *(Solidago canadensis)* positioned in the same arrangement in each mesocosm. More details regarding the establishment of mesocosms and climate-extreme treatments are provided in ref et al.

Drought and warming treatments were applied annually beginning in 2023, and the same treatment combinations were maintained across years. Drought was simulated using rain-out shelters placed over each mesocosm during the growing season, beginning in late May and remaining in place for approximately 1.5 months until late June each year (Fig. 1). At the end of the drought period, soil moisture in the upper 0–10 cm declined to ≤5% volumetric water content in mesocosm treated with drought, indicating extreme drought conditions as defined by the U.S. Drought Monitor (National Drought Mitigation Center, USDA, and NOOA). Warming treatments were applied using infrared heaters during the growing season (mid-March to late October), and consisted of four warming regimes: (1) ‘Ambient’ (ambient temperature conditions), (2) ‘Constant warming’ (W) in which plots were constantly heated at 2°C throughout the growing season, (3) ‘Heat waves’ (HW) with pulses of plus 7°C for 1 week each month (seven times per year), and (4) ‘Constant warming and heat waves’ (W + HW). The ‘Constant warming’ treatment simulates the projected increase in global mean temperature under moderate climate warming scenarios (IPCC, 2023). Heat wave treatment was designed to simulate periodic extreme warming events projected to increase in intensity and frequency under climate change (Marx et al., 2021; IPCC, 2023). The warming regimes were fully crossed with two moisture regimes, ambient precipitation and drought, with five replicate mesocosms per treatment combination, resulting in 40 mesocosms (4 warming regimes × 2 moisture regimes × 5 replicates; Fig. 1).

Soil samples for this study were collected at three time points in 2024 and 2025 (Fig. 1). Therefore, samples collected before annual drought manipulation captured potential legacy effects of drought and warming treatments applied during previous years, allowing us to assess the accumulated effects of drought. Here, we define the ’drought legacy’ as the persistent effects of drought on microbial responses that were detectable prior to subsequent annual drought manipulation (Canarini et al., 2021; Evans et al., 2022). Each year, soils were sampled during three campaigns: 1) Drought legacy, before annual drought onset (mid-May), 2) Resistance phase, at the end of drought (late June), 3) Recovery phase, one month after rewetting (late July) (Fig. 1). Soil composite samples were obtained by mixing two soil cores (2.5 cm in diameter, 0-10 cm in depth) randomly collected within a designated quadrant in each mesocosm. The mesocosms were divided into four quadrants, and the sampled quadrant was rotated among sampling campaigns to avoid resampling disturbed soil. The collected soils were sieved through a 2 mm sieve, stored at 4°C immediately after collection, and used for microbial activity and metabolic functions assays within one week.

### 2.2 Assays

#### 2.2.1 Microbial extracellular enzyme activities

The acquisition of potential microbial resources was evaluated for 2 years (2024 and 2025) (Table S1) by measuring the potential activity of six key extracellular enzymes involved in the cycling of carbon (C), nitrogen (N) and phosphorus (P) (Table S2). Soil slurries were prepared by mixing 2 g of fresh soil with 60 mL of a Tris-buffer solution (0.1 M, pH 8.0), appropriate for neutral to alkaline soil conditions, and homogenized with a magnetic stirrer. Aliquots (200 µL) of the slurry were dispensed into a 96-well microplate with 50 µL of MUB-linked substrates (4-methylumbelliferone) for β-glucosidase (BG), N-acetylglucosaminidase (NAG), and phosphatase (PHO), and incubated in the dark at 25°C for 2 h (BG, NAG) or 3 h (PHO) (Table S2). Leucine aminopeptidase (LAP) was measured by combining soil slurry (200 µL) with 50 µL of substrate linked to AMC (L-leucine-7-amido-4-methylcoumarin) and incubated in the dark at 25°C for 3 h. Extracellular enzymatic assays were incubated at 25°C to assess potential microbial activity within the near optimal range for many mesophilic soil microorganisms. Fluorescence was measured at excitation/emission wavelengths of 365/445 nm for MUB and 365/450 nm for AMC using a BioTek Synergy H1 microplate reader (BioTek Instruments Inc., USA).

Phenol oxidase (POX) and peroxidase (PER) were measured by mixing 400 µL of soil slurry with 400 µL of 3,4-dihydroxy-L-phenylalanine (L-DOPA). For the determination of peroxidase, 10 µL of 0.3% H_2_O_2_ was added to the designated wells, while the wells without H_2_O_2_ were used to quantify the activity of phenol oxidase. Activities were quantified spectrophotometrically by measuring the absorbance of oxidized L-DOPA at 460 nm, immediately after plate preparation and after incubation (Sinsabaugh, 2010). The plates were incubated in the dark at 25°C, PER for 5 h and POX for 1 h. The resource acquisition enzymes (BG, NAG, PHO, LAP, POX, and PER) were expressed as µmol product formed g^-1^ dry soil h⁻¹.

#### 2.2.2 Dehydrogenase activity

As a proxy for potential active oxidative metabolism, soil dehydrogenase activity (DEH) was measured for 1 year (2025) (Fig. 1, Table S2) following Von Mersi & Schinner, (1991) with minor modifications (Camiña et al., 1998). Briefly, 1.0 g of oven dry equivalent of field moist soil was mixed with 1.5 mL of Tris buffer (pH = 7.5) and 2.0 mL of iodonitrotetrazolium chloride (INT). The soil samples were sealed and incubated in the dark and under constant mixing at 40°C for 2 h, following the standard INT reduction protocol. Following incubation, 5 mL of N, N-dimethylformamide (DMF): Ethanol extractant (1: 1) was added to each tube and the samples were shaken for 1 h at 25°C in the dark. The suspensions were centrifuged (3,900 x g for 10 minutes) and 2 mL of supernatant was filtered through a 0.45 µm PTFE syringe filter, before measuring absorbance at 490 nm in 96-well plates (300 µl). A calibration curve was prepared using iodonitrotetrazolium formazan (INTF) standards (1 mM). Dehydrogenase was expressed as µmol INTF g^-1^ dry soil h^-1^.

#### 2.2.3 Microbial glycogen pools

As a proxy for microbial C storage, microbial glycogen (GLY) was measured for 1 year (2025) (Fig.1, Table S2). Before glycogen quantification, microbial cells were recovered from the soil using cell-extraction procedure based on (Lindahl & Bakken, 1995) with modification. Briefly, 2 g of fresh soil were suspended in 10 mL of sterile ice-cold sodium pyrophosphate solution, 0.2% w/v, adjusted to pH 8.5. The soil suspension was shaken reciprocally at 125 rpm for 2 h at 4°C to promote physical dislodgment of microbial cells from the soil matrix. The suspension was then centrifuged at 200 x g for 5 min at 4°C to sediment coarse soil particles and debris. For density-based separation of microbial cells, 6 mL of Nycodenz solution, density 1.3 g mL ^-1^, was layered beneath the homogenized soil suspension. The samples were centrifuged at 10,000 x g for 30 min at 4°C. The microbial cell-enriched fraction was recovered from the layer above the Nycodenz cushion, washed with 0.85% NaCl, and centrifuged (15,000 x g, 10 min) to obtain cell pellets. Glycogen was extracted in 30% KOH at 95°C for 60 min, precipitated with 1 mL of ice-cold 100% ethanol, and incubated at -20°C for 30 min (Zavřel et al., 2018; Vidal & Venegas-Calerón, 2019). The samples were centrifuged twice (20,000 x g for 10 min at 4°C), with an intermediate wash in 70% ethanol (500 µL). The dried pellet was hydrolyzed with 100 µL 1 N HCl at 95°C for 15 min, neutralized with 100 µL 1N NaOH, and diluted with 150 µL of Milli-Q water. The samples (300 µL) were mixed with 1 mL of anthrone reagent, incubated at 95°C for 10 min, and the absorbance was measured at 620 nm using a BioTek Synergy H1 microplate reader (BioTek Instruments Inc., USA). Glycogen was expressed as µmol glucose equivalents g^-1^ dry soil.

#### 2.2.4 Enzymatic stoichiometry

To assess potential microbial nutrient acquisition strategies, we calculated enzymatic stoichiometric ratios following Sinsabaugh et al. (2009), using activities of β-glucosidase (BG) as a proxy for carbon acquisition, N-acetylglucosaminidase (NAG) and leucine aminopeptidase (LAP) for nitrogen acquisition, and phosphatase (PHO) for phosphorus acquisition (Sinsabaugh et al., 2009). We also assessed the ratio of oxidases to hydrolases (Ox: Hy) as an index of relative investment in enzymes targeting chemically complex versus labile C substrates, with higher ratios indicating relatively greater investment in the decomposition of chemically recalcitrant C pools (Romero-Olivares et al., 2017; Chen et al., 2020). Ox represents oxidative C-acquisition, quantified as the combined activity of phenol oxidase (POX) and peroxidase (PER), whereas Hy represents hydrolytic C acquisition, quantified as β-glucosidase (BG) activity.

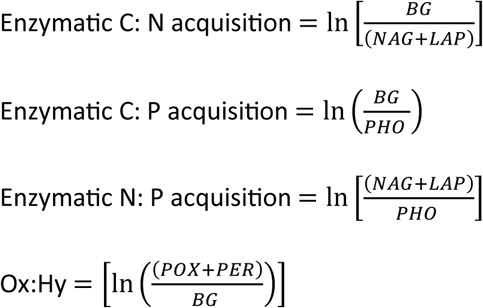

### 2.3 Statistical analyses

All analyses were conducted in R version 4.5.0 (R Core Team, 2025). To quantify the effect of drought across warming regimes on enzymatic activities and ratios, dehydrogenase activity, and glycogen pools, we fit linear mixed-effects models (LMMs) using the *glmmTMB* function of the glmmTMB package (v 1.1.8) (Brooks et al., 2017). Extracellular enzyme activities were log-transformed, whereas dehydrogenase activity and glycogen pools were analysed without transformation. Enzymatic stoichiometric ratios were calculated as natural log-transformed ratios and analysed without further transformation. The models included drought, warming regime, sampling phase, and their interactions as fixed effects, with plot (experimental unit) as a random effect (Brooks et al., 2017). The significance of fixed effects was evaluated using Type II Wald χ² tests. Extracellular enzyme activities were measured for 2 years (2024 and 2025; n = 10 per treatment), while dehydrogenase activity (DEH), glycogen (GLY) and log response ratios (LRR) used data of 2025 (n = 5 per treatment). Accordingly, Year was included as an additional random effect only for enzymatic activities and ratios (Fig.1 and Table S2) (Brooks et al., 2017). Furthermore, we fit a joint model with the log response ratios (LRRs) of microbial extracellular C-acquisition (Hy and Ox), and intracellular C storage (GLY), specifying an unstructured covariance matrix. Model diagnostics were assessed using simulated residuals with the DHARMa package (v 0.4.7), including tests for uniformity, dispersion, outliers, and residual patterns against fitted values and predictors (v 0.4.7) (Hartig, 2016). Estimated marginal means were computed using the emmeans package (v 1.11.2) (Lenth, 2025). Effect sizes (Cohen’s d) were calculated from the estimated mean comparison between drought and ambient precipitation treatments (drought – control) within each warming regime and sampling phase in each year, and were calculated with the *eff_size* function of our mixed-effects models (Lenth, 2025). For visualization, we used the ggplot2 package (v 3.5.2) (Wickham, 2016).

## 3. Results

### 3.1 Drought effects on potential resource acquisition, potential active oxidative metabolism, and internal C storage across warming regimes

Drought and warming legacy effects were detected in various resource acquiring enzymes. During the drought legacy phase, drought reduced the activity of BG by 9.9% under constant warming + heat waves (‘W + HW’), while the activity of leucine aminopeptidase (LAP) was reduced by 41.7% under constant warming (‘W’) (Fig. 2, Table S3, Figs. S1-2). In contrast, PHO activity increased by 55.4% under constant warming + heat waves (‘W + HW’) (Fig. 2, Table S3, Figs. S1-2). No significant responses were detected for the enzymatic stoichiometric ratios (Fig. 3, Table S4, Fig. S3). Drought also reduced potential active oxidative metabolism (DEH) and C storage pools (GLY) by 25.3% and 58.4%, respectively. No effects of drought x warming interaction were observed for DEH activity or GLY in the legacy phase (Fig. 4, Table S5, Fig. S4).

**Figure 2.**
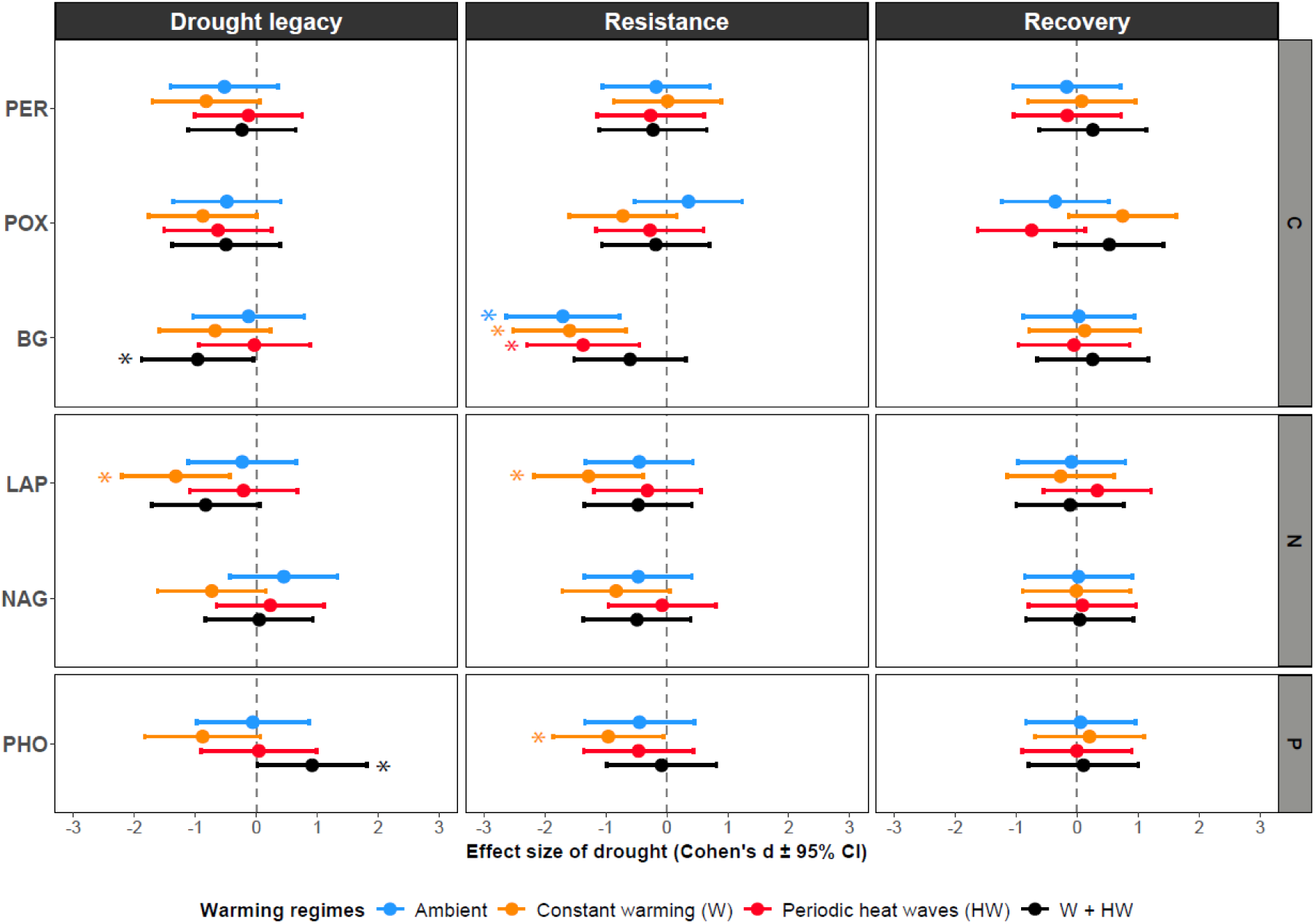
Effects of drought on microbial potential resource acquisition across warming regimes. Effect sizes (Cohen’s d ± 95% CI) represent the standardized difference in enzyme activity between drought and control plots under four warming treatments, measured at three time points: Drought legacy (before drought), Resistance (at the end of the drought), and Recovery (one month after rewetting). Panels are displayed by nutrient-association groups (C-, N-, or P-acquiring enzymes). PER = peroxidase; 334 POX = phenol oxidase; BG = β-glucosidase; LAP = leucine aminopeptidase; NAG = N acetylglucosaminidase; PHO = phosphatase. Asterisks indicate significant effects (95% CI not overlapping with zero).

**Figure 3.**
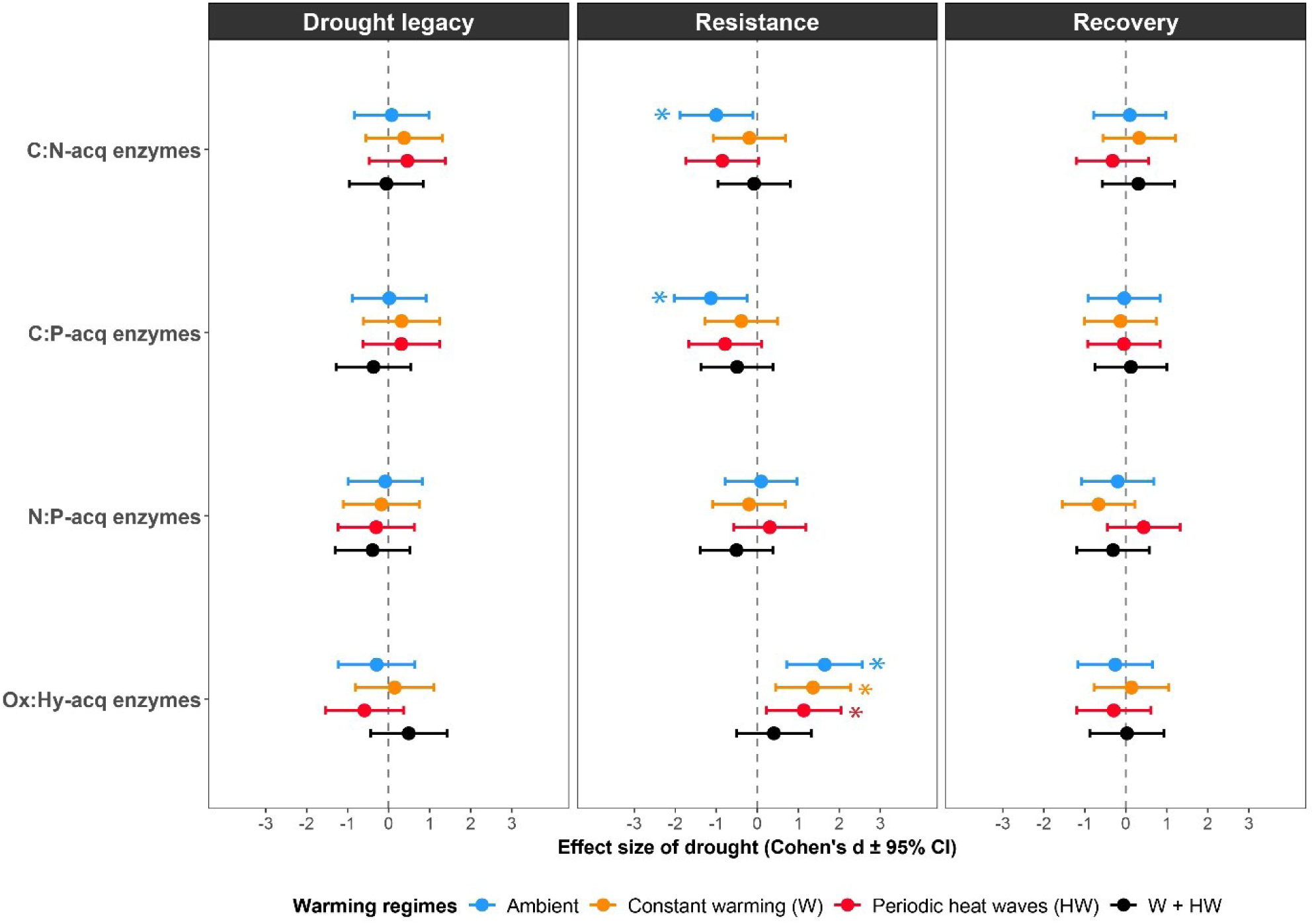
Effects of drought on potential resource acquisition enzymatic ratios across warming regimes. Effect sizes (Cohen’s d ± 95% CI) represent the standardized difference in enzymatic ratios between drought and control plots under four warming treatments, measured at three time points: Drought legacy (before drought), Resistance (at the end of the drought), and Recovery (one month after rewetting). Enzymatic ratios are shown as follows: C: N = ln (BG/(NAG+LAP)); N: P = ln ((NAG+LAP)/PHO)); C: P = ln (BG/PHO); Oxidases: Hydrolases = ln (POX+PER/ BG). Asterisks indicate significant effects. Asterisks indicate significant effects (95% CI not overlapping with zero).

**Figure 4.**
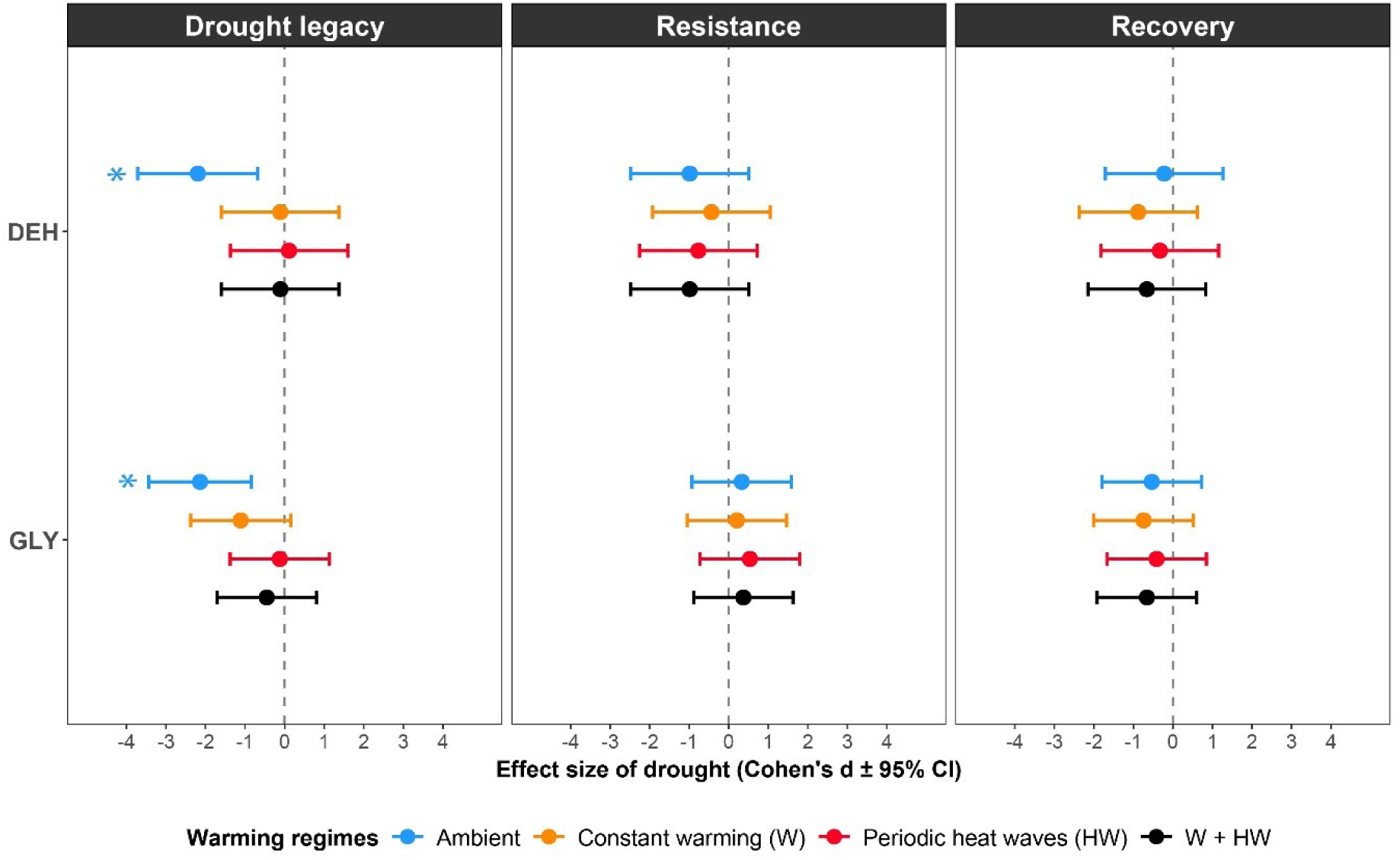
Effects of drought on potential microbial active oxidative metabolism and internal C storage across warming regimes. Effect sizes (Cohen’s d ± 95% CI) represent the standardized effect of drought on potential microbial active oxidative metabolism and internal C storage across four warming regimes. GLY = glycogen, DEH = dehydrogenase, measured at three time points: Drought legacy (before drought), Resistance (at the end of the drought), and Recovery (one month after rewetting) (n= 5). Asterisks indicate significant effects (95% CI not overlapping with zero).

During the resistance phase, drought reduced BG activity by 48.1% under ambient temperature conditions, by 45.9% under constant warming (‘W’), and by 41.0% under heat waves relative to control conditions (Fig. 2, Table S3, Fig. S1). LAP and PHO activities were reduced only when drought was combined with constant warming (‘W’), by 40.8% and 37.0%, respectively (Fig. 2, Table S3, Fig. S1). On the contrary, the oxidases (PER and POX) did not respond significantly to drought, warming, or their interaction (Fig. 2, Table S3, Fig. S1). Drought reduced the enzymatic C: N and C: P ratios by 37.5% and 35.7% under ambient temperature conditions, respectively, with no significant drought x warming interaction. In contrast, the ratio of oxidases to hydrolases (Ox: Hy) increased by 92.1% under drought, 71.7% under drought and constant warming (‘W’), and 56.9% under drought and heat waves (‘HW’) (Fig. 3, Table S4. Fig. S3). Consistent with this change in the balance between extracellular C acquisition strategies, joint analysis of log response ratios (LRRs) revealed a significant three-way interaction among microbial C processes (Hy, Ox, and GLY), climate extreme treatments, and sampling phase (C proxy x Extreme x Sampling: χ^2^ = 55.00, df = 24, p<0.001), indicating that the relative responses of these C processes differed across sampling phases (Table S6-7). Significant interactions of the C proxy × Extreme (χ² = 32.70, df = 12, P = 0.001) and the C proxy × Sampling (χ² = 84.13, df = 4, P < 0.001) interactions further supported these results (Table S6 and S7). The N: P ratio did not show a significant response under drought or under the drought x warming interaction effects (Fig. 3, Table S4). Furthermore, we did not observe significant drought effects or drought × warming interactions on DEH activity or GLY pools (Fig. 4, Table S5).

During the recovery phase (one month after rewetting), extracellular enzyme activities did not differ between drought and control plots between warming regimes, indicating recovery of all extracellular enzyme activities (Fig. 2, Table S3, Figs. S1-S2). Enzymatic stoichiometric ratios also returned to control levels during the recovery phase (Fig. 3, Table S4, Fig. S3). Finally, active microbial oxidative metabolism and internal C storage pools did not show any treatment difference and no significant drought x warming interaction was found in the recovery phase (Fig. 4, Table S5, Fig. S4).

## 4. Discussion

Our results showed that drought primarily altered microbial internal C storage during the legacy phase and reduced microbial potential resource acquisition during the resistance phase, while potential active oxidative metabolism and microbial internal C storage remained comparatively stable across warming regimes. Following rewetting, all measured proxies of resource acquisition and storage fully recovered. Our first hypothesis was partly supported, drought reduced potential resource acquisition while maintaining potential oxidative metabolism during the resistance phase. However, contrary to our expectations, microbial internal C storage did not increase during the resistance phase, and the predicted under-recovery of resource acquiring enzymes after rewetting was not observed. Contrary to our second hypothesis, neither constant warming nor heat waves amplified drought constraints in specific extracellular enzymes during the resistance phase or induced over-recovery of measured proxies. However, both warming regimes increased Oxidases: Hydrolases ratio during the resistance phase, consistent with our expectations. In addition, drought effects in combination with warming extended to N-and P-acquiring enzymes (Fig. 2). Contrary to our third hypothesis, drought combined with constant warming and heat waves did not impose the strongest functional constraints either in under-recovery following rewetting in any measured proxy. Together, these findings suggest that soil microorganisms shifted resource allocation under climate extremes, and these shifts were phase dependent. Drought legacy effects were most pronounced in microbial internal C storage, while during peak drought soil resource acquisition was mainly reduced while maintaining potential oxidative metabolism and internal C storage. After rewetting, these functional activities recovered, suggesting that these physiological and metabolic adjustments could be microbial survival strategies under variable climatic conditions.

Drought legacy effects were function-specific and strongest for glycogen concentration, whereas it remained stable during the subsequent resistance and recovery phase (Fig. 4). This phase-dependent response suggests that microbial C storage was sensitive to previous drying-rewetting cycles but resistant to the subsequent drought event, consistent with the mobilization of intracellular C reserves after drought. Joint analysis of extracellular C acquisition and internal C storage also indicated greater hydrolytic C-acquiring enzyme activity relative to glycogen under drought x heat waves (Table S6), consistent with a shift from intracellular C storage to extracellular resource acquisition. However, because these responses were not detected in the individual models (Figs. 2 and 4), this interpretation should be treated with caution. Additionally, plant-mediated pathways may have contributed to these legacy effects. In a previous study using the same mesocosm experiment, drought legacies were detected in plant biomass and functional traits (Tartini et al., 2026). Drought-induced changes in root exudation could therefore have reduced the supply of labile C to soil microorganisms and altered the microbial, as proposed in previous studies (De Vries et al., 2012; Canarini et al., 2021). Alternatively, persistent changes in microbial community composition may have contributed to the observed response. In particular, drought may have selected fast growing and rapidly recovering taxa that may explain the rapid recovery of microbial functions despite persistent legacy effects (Evans & Wallenstein, 2014; De Nijs et al., 2019; Oram et al., 2025). However, because our measurements coincided with the early growing season, drought legacy cannot be fully separated from concurrent plant responses.

During the resistance phase, the potential activities of extracellular enzymes declined, while the activity of dehydrogenase and the concentration of glycogen remained stable (Figs. 2-4). This decoupling suggests that drought constrained extracellular resource acquisition more strongly than potential intracellular oxidative metabolism or microbial C storage. By restricting the diffusion of enzymes, substrates, and reaction products, low soil moisture may reduce the return on metabolically costly production of extracellular enzymes (Malik et al., 2020; Fanin et al., 2022). Oxidase activity remained stable, consistent with previous drought x warming studies (e.g., Yan et al., 2020), but drought increased the C Oxidases: Hydrolases ratio and our joint analysis further revealed lower BG relative to oxidases (Table S6-7). Because absolute oxidase activity did not increase, these results indicate a relative shift toward oxidative C acquisition driven primarily by suppression of hydrolytic activity. This pattern is consistent with the reduced accessibility of labile C under drought (Romero-Olivares et al., 2017; Chen et al., 2020; Sinsabaugh, 2010). Contrary to our expectations, the glycogen concentration did not change during drought, either alone or in combination with warming. By contrast, Canarini et al. (2024) reported a five-fold increase in microbial triglycerides under drought. Although both compounds function as intracellular C storage compounds, glycogen is a readily mobilizable carbohydrate reserve, while triglycerides represent long-term lipid storage (Mason-Jones et al., 2022). Therefore, differences in their biochemical functions may contribute to their contrasting responses, although variation in experimental conditions and microbial communities between studies may also be important.

One month after rewetting, extracellular enzyme activities, enzymatic ratios, dehydrogenase activity, and glycogen concentration did not differ between control and drought plots across warming regimes. This is likely because rewetting restored aqueous diffusion and substrate accessibility. Similarly, Cordero et al. (2023) observed complete recovery under milder drought, but an under-recovery of extracellular enzyme activities under intense drought even after 6 months of rewetting, which they attributed to reduced soil C availability. In our experiment, the custom-formulated substrate contained 10% garden compost, which may have increased labile C availability and facilitated recovery after diffusion and substrate availability were restored after rewetting. Nevertheless, repeated climatic extremes may progressively alter substrate availability, with longer-term consequences for soil C storage and ecosystem functioning (Allison, 2023).

Finally, the asymmetric responses of glycogen and extracellular enzymes between drought phases (Table S6-7) suggest a temporal shift in the allocation of microbial resources, suggesting intracellular reserves mobilization after drying-rewetting cycles but maintenance during peak drought. These shifts during different drought phases are consistent with physiological acclimatization strategies that prioritize survival over growth under drying-rewetting cycles (Schimel et al., 2007; Leizeaga et al., 2021). Previous work has shown that repeated drought can alter microbial C use without reducing microbial resistance or resilience, potentially reflecting physiological adaptations to drying-rewetting cycles or indirect effects through altered substrate availability and plant productivity (Leizeaga et al., 2021). However, these responses might also reflect variation in microbial biomass or substrate availability, which was not measured in this study. Standardizing them by microbial biomass would help distinguish changes in microbial abundance from biomass-specific activity (Piton et al., 2020). Furthermore, extracellular enzyme activities, dehydrogenase activity, and glycogen concentrations were measured under standardized laboratory conditions and should therefore be interpreted as potential responses of a grassland model system rather than process rates in situ of natural systems. Moreover, integrating these approaches with isotope measurements of microbial growth and carbon use efficiency, together with microbial community composition and stress tolerance mechanisms, would give insights into microbial physiological adjustment under stress (Malik et al., 2020; Canarini et al., 2021). Finally, our results suggest that microbial physiological shifts are phase dependent and may function as microbial survival strategies under compound extremes, possibly contributing to fast recovery following rewetting despite drought legacies (Mason-Jones et al., 2022; Allison, 2023; Canarini et al., 2024).

## 4. Conclusions

Overall, our study shows that drought was the primary driver of measured microbial functional responses, while warming and heat waves modulated these drought effects depending on the functional activity and the sampling phase. During the legacy phase, drought had the strongest effects on microbial internal C storage, while under warming these legacy effects also extended to potential extracellular enzyme activities. Drought reduced labile C acquisition, and under warming its effects extended to other acquisition enzymes. The microbial communities maintained active oxidative metabolism, while the enzyme ratios suggested a relative shift toward enzymes associated with complex C substrates and lower extracellular acquisition relative to glycogen pools, consistent with shifts in microbial functional allocation. Despite these shifts during the drought legacy and resistance phases, the functional proxies returned to control levels one month after rewetting, indicating potential functional resilience in soil microorganisms. These responses may reflect physiological shifts that function as microbial survival strategies under compound extremes, and may contribute to fast recovery after rewetting (Mason-Jones et al., 2022; Allison, 2023; Canarini et al., 2024). However, repeated drying-rewetting cycles under warming might alter the balance between microbial extracellular resource acquisition and internal C storage, and we still don’t know if it might generate accumulated microbial physiological or metabolic costs associated with stress tolerance (Martínez-De León & Thakur, 2024), with consequences on microbial recovery trajectories (Thakur et al., 2022), and ultimately on ecosystem functions (Allison, 2023). Overall, our findings show that climate extremes alter microbial allocation strategies during drought and its legacy, while maintaining rapid functional recovery after rewetting, suggesting that these physiological adjustments may function as survival strategies under increasingly variable climates.

