## Supplementary Table 1-6, Figures 1-4 for "Resource acquisition is more sensitive than carbon storage in soil microorganisms under climate extremes"

**Supplementary Table 1.** Potential ecological proxies, analytical proxies, and years of measurements used for analysis.

| Potential microbial functional activity and storage | Analytical proxy | Years used for analysis |
|---|---|---|
| **Resource acquisition** | Extracellular enzymes (EEA) | 2024 and 2025 |
| **Internal Carbon (C) storage** | Glycogen (GLY) | 2025 |
| **Active oxidative metabolism** | Dehydrogenase activity (DEH) | 2025 |

**Supplementary Table 2.** Overview and characteristics of the microbial polymers, extracellular and intracellular enzymes.

| Enzyme / polymer | Abbreviation | Function | Class | Activity | Reference |
|---|---|---|---|---|---|
| **Phenol oxidase** | POX | Nutrient acquisition (Extracellular enzymes) | Oxidases | Oxidizes phenolic compounds, key in lignin and humus degradation. | (Sinsabaugh, 2009) |
| **Peroxidase** | PER | | | Breaks down complex aromatic polymers using $H_2O_2$; lignin degradation. | |
| **β-Glucosidase** | BG | | Hydrolases | Hydrolyses cellobiose and other short cellulose-derived oligosaccharides into glucose, completing cellulose decomposition and contributing to C cycling. | |
| **Leucine aminopeptidase** | LAP | | | Cleaves N-terminal leucine residues from peptides, releasing amino acids and contributing to microbial N-acquisition. | |
| **β-N-Acetylglucosaminidase** | NAG | | | Degrades chitin & peptidoglycan, releases N & C. | |
| **Acid phosphatase** | PHO | | | Mineralizes organic P to release inorganic phosphate. | |
| **Glycogen** | GLY | Storage (polymer) | - | Microbial intracellular carbon storage compound (energy reserve). Present in bacteria, protozoa, fungi and animal cells. | (Vidal & Venegas-Caleron, 2019; Mason-Jones, 2021) |
| **Dehydrogenase** | DEH | Intracellular enzyme (cytoplasmic) | Oxidoreductase | Intracellular enzyme. Key enzyme for oxidation processes such as respiration. | (Malachowska-Jutsz, 2019). |

**Supplementary Table 3.** Output of the mixed-effect model to quantify the effects of drought on nutrient acquisition enzymes across warming regimes at 3 different sampling points. A model was run per each enzyme, it included an interaction effect of Drought (D), Treatment (warming regimes), and Sampling point (Sampling 1 = Drought legacy, Sampling 2 = Resistance, Sampling 3 = Recovery). It counted for spatial and yearly variation by including Plot and Year (2024 and 2025) as random effects. Significant effects are marked in bold. SE stands for standard error.

| Nutrient acquisition enzyme | Predictor | Estimate | SE | Z-value | p-value |
|---|---|---|---|---|---|
| **POX** | (Intercept) | 1.488 | 0.246 | 6.048 | <0.001 |
| | D | 0.102 | 0.094 | 1.079 | 0.280 |
| | TreatmentAmbient + 2C | -0.049 | 0.094 | -0.518 | 0.604 |
| | TreatmentAmbient + Periodic Heat Wave | -0.037 | 0.094 | -0.391 | 0.696 |
| | TreatmentAmbient + 2C + Periodic Heat Wave | -0.063 | 0.094 | -0.665 | 0.506 |
| | Sampling2 | 0.061 | 0.094 | 0.650 | 0.516 |
| | **Sampling3** | **0.194** | **0.094** | **2.057** | **0.040** |
| | D:TreatmentAmbient + 2C | 0.083 | 0.133 | 0.623 | 0.533 |
| | D:TreatmentAmbient + Periodic Heat Wave | 0.031 | 0.133 | 0.232 | 0.817 |
| | D:TreatmentAmbient + 2C + Periodic Heat Wave | 0.003 | 0.133 | 0.021 | 0.983 |
| | D:Sampling2 | -0.177 | 0.133 | -1.328 | 0.184 |
| | D:Sampling3 | -0.026 | 0.133 | -0.196 | 0.844 |
| | TreatmentAmbient + 2C:Sampling2 | -0.143 | 0.133 | -1.076 | 0.282 |
| | TreatmentAmbient + Periodic Heat Wave:Sampling2 | -0.074 | 0.133 | -0.557 | 0.578 |
| | TreatmentAmbient + 2C + Periodic Heat Wave:Sampling2 | 0.022 | 0.133 | 0.162 | 0.872 |
| | TreatmentAmbient + 2C:Sampling3 | 0.109 | 0.133 | 0.814 | 0.416 |
| | TreatmentAmbient + Periodic Heat Wave:Sampling3 | -0.029 | 0.133 | -0.215 | 0.830 |
| | TreatmentAmbient + 2C + Periodic Heat Wave:Sampling3 | 0.172 | 0.133 | 1.291 | 0.197 |
| | D:TreatmentAmbient + 2C:Sampling2 | 0.144 | 0.188 | 0.761 | 0.446 |
| | D:TreatmentAmbient + Periodic Heat Wave:Sampling2 | 0.102 | 0.188 | 0.541 | 0.588 |
| | D:TreatmentAmbient + 2C + Periodic Heat Wave:Sampling2 | 0.110 | 0.188 | 0.585 | 0.558 |
| | D:TreatmentAmbient + 2C:Sampling3 | -0.316 | 0.188 | -1.676 | 0.094 |
| | D:TreatmentAmbient + Periodic Heat Wave:Sampling3 | 0.051 | 0.188 | 0.269 | 0.788 |
| | D:TreatmentAmbient + 2C + Periodic Heat Wave:Sampling3 | -0.189 | 0.188 | -1.005 | 0.315 |
| **PER** | (Intercept) | 0.715 | 0.164 | 4.354 | <0.001 |
| | D | 0.227 | 0.195 | 1.168 | 0.243 |
| | TreatmentAmbient + 2C | -0.267 | 0.195 | -1.374 | 0.169 |
| | TreatmentAmbient + Periodic Heat Wave | -0.038 | 0.195 | -0.195 | 0.845 |
| | TreatmentAmbient + 2C + Periodic Heat Wave | -0.062 | 0.195 | -0.320 | 0.749 |
| | **Sampling2** | **0.494** | **0.195** | **2.538** | **0.011** |
| | Sampling3 | 0.161 | 0.195 | 0.830 | 0.406 |
| | D:TreatmentAmbient + 2C | 0.130 | 0.275 | 0.474 | 0.635 |
| | D:TreatmentAmbient + Periodic Heat Wave | -0.171 | 0.275 | -0.623 | 0.533 |
| | D:TreatmentAmbient + 2C + Periodic Heat Wave | -0.125 | 0.275 | -0.454 | 0.650 |
| | D:Sampling2 | -0.152 | 0.275 | -0.553 | 0.580 |
| | D:Sampling3 | -0.153 | 0.275 | -0.556 | 0.578 |
| | TreatmentAmbient + 2C:Sampling2 | 0.248 | 0.275 | 0.902 | 0.367 |
| | TreatmentAmbient + Periodic Heat Wave:Sampling2 | 0.058 | 0.275 | 0.210 | 0.834 |
| | TreatmentAmbient + 2C + Periodic Heat Wave:Sampling2 | 0.086 | 0.275 | 0.312 | 0.755 |

| | | | | | |
|---|---|---|---|---|---|
| | TreatmentAmbient + 2C:Sampling3 | 0.333 | 0.275 | 1.210 | 0.226 |
| | TreatmentAmbient + Periodic Heat Wave:Sampling3 | 0.036 | 0.275 | 0.132 | 0.895 |
| | TreatmentAmbient + 2C + Periodic Heat Wave:Sampling3 | 0.266 | 0.275 | 0.965 | 0.334 |
| | D:TreatmentAmbient + 2C:Sampling2 | -0.212 | 0.389 | -0.544 | 0.587 |
| | D:TreatmentAmbient + Periodic Heat Wave:Sampling2 | 0.211 | 0.389 | 0.543 | 0.587 |
| | D:TreatmentAmbient + 2C + Periodic Heat Wave:Sampling2 | 0.147 | 0.389 | 0.378 | 0.706 |
| | D:TreatmentAmbient + 2C:Sampling3 | -0.237 | 0.389 | -0.609 | 0.542 |
| | D:TreatmentAmbient + Periodic Heat Wave:Sampling3 | 0.168 | 0.389 | 0.431 | 0.667 |
| | D:TreatmentAmbient + 2C + Periodic Heat Wave:Sampling3 | -0.061 | 0.389 | -0.158 | 0.875 |
| **BG** | (Intercept) | 4.619 | 0.259 | 17.837 | <0.001 |
| | D | 0.049 | 0.178 | 0.274 | 0.784 |
| | TreatmentAmbient + 2C | -0.093 | 0.178 | -0.523 | 0.601 |
| | **TreatmentAmbient + Periodic Heat Wave** | **0.473** | **0.178** | **2.651** | **0.008** |
| | TreatmentAmbient + 2C + Periodic Heat Wave | 0.104 | 0.178 | 0.582 | 0.560 |
| | Sampling2 | -0.315 | 0.172 | -1.828 | 0.068 |
| | **Sampling3** | **0.459** | **0.172** | **2.665** | **0.008** |
| | D:TreatmentAmbient + 2C | 0.211 | 0.252 | 0.838 | 0.402 |
| | D:TreatmentAmbient + Periodic Heat Wave | -0.036 | 0.252 | -0.144 | 0.886 |
| | D:TreatmentAmbient + 2C + Periodic Heat Wave | 0.322 | 0.252 | 1.277 | 0.202 |
| | **D:Sampling2** | **0.607** | **0.244** | **2.491** | **0.013** |
| | D:Sampling3 | -0.060 | 0.244 | -0.246 | 0.806 |
| | TreatmentAmbient + 2C:Sampling2 | 0.206 | 0.244 | 0.847 | 0.397 |
| | TreatmentAmbient + Periodic Heat Wave:Sampling2 | -0.444 | 0.244 | -1.823 | 0.068 |
| | TreatmentAmbient + 2C + Periodic Heat Wave:Sampling2 | 0.185 | 0.244 | 0.758 | 0.448 |
| | TreatmentAmbient + 2C:Sampling3 | -0.035 | 0.244 | -0.142 | 0.887 |
| | **TreatmentAmbient + Periodic Heat Wave:Sampling3** | **-0.509** | **0.244** | **-2.089** | **0.037** |
| | TreatmentAmbient + 2C + Periodic Heat Wave:Sampling3 | 0.091 | 0.244 | 0.374 | 0.708 |
| | D:TreatmentAmbient + 2C:Sampling2 | -0.254 | 0.345 | -0.736 | 0.461 |
| | D:TreatmentAmbient + Periodic Heat Wave:Sampling2 | -0.092 | 0.345 | -0.266 | 0.791 |
| | **D:TreatmentAmbient + 2C + Periodic Heat Wave:Sampling2** | **-0.746** | **0.345** | **-2.165** | **0.030** |
| | D:TreatmentAmbient + 2C:Sampling3 | -0.248 | 0.345 | -0.719 | 0.472 |
| | D:TreatmentAmbient + Periodic Heat Wave:Sampling3 | 0.067 | 0.345 | 0.196 | 0.845 |
| | D:TreatmentAmbient + 2C + Periodic Heat Wave:Sampling3 | -0.409 | 0.345 | -1.186 | 0.236 |
| **NAG** | (Intercept) | 2.858 | 0.210 | 13.585 | <0.001 |
| | D | -0.283 | 0.280 | -1.009 | 0.313 |
| | TreatmentAmbient + 2C | -0.494 | 0.280 | -1.761 | 0.078 |
| | TreatmentAmbient + Periodic Heat Wave | 0.341 | 0.280 | 1.216 | 0.224 |
| | TreatmentAmbient + 2C + Periodic Heat Wave | -0.375 | 0.280 | -1.338 | 0.181 |
| | **Sampling2** | **-0.554** | **0.280** | **-1.977** | **0.048** |
| | Sampling3 | -0.158 | 0.280 | -0.563 | 0.574 |
| | D:TreatmentAmbient + 2C | 0.741 | 0.397 | 1.868 | 0.062 |
| | D:TreatmentAmbient + Periodic Heat Wave | 0.139 | 0.397 | 0.351 | 0.726 |
| | D:TreatmentAmbient + 2C + Periodic Heat Wave | 0.253 | 0.397 | 0.637 | 0.524 |
| | D:Sampling2 | 0.576 | 0.397 | 1.453 | 0.146 |
| | D:Sampling3 | 0.270 | 0.397 | 0.680 | 0.497 |

| | | | | | |
|---|---|---|---|---|---|
| | TreatmentAmbient + 2C:Sampling2 | 0.341 | 0.397 | 0.861 | 0.389 |
| | TreatmentAmbient + Periodic Heat Wave:Sampling2 | -0.316 | 0.397 | -0.798 | 0.425 |
| | TreatmentAmbient + 2C + Periodic Heat Wave:Sampling2 | 0.488 | 0.397 | 1.232 | 0.218 |
| | TreatmentAmbient + 2C:Sampling3 | 0.270 | 0.397 | 0.681 | 0.496 |
| | TreatmentAmbient + Periodic Heat Wave:Sampling3 | -0.432 | 0.397 | -1.089 | 0.276 |
| | TreatmentAmbient + 2C + Periodic Heat Wave:Sampling3 | 0.286 | 0.397 | 0.720 | 0.472 |
| | D:TreatmentAmbient + 2C:Sampling2 | -0.514 | 0.561 | -0.916 | 0.360 |
| | DTreatmentAmbient + Periodic Heat Wave:Sampling2 | -0.385 | 0.561 | -0.686 | 0.492 |
| | D:TreatmentAmbient + 2C + Periodic Heat Wave:Sampling2 | -0.240 | 0.561 | -0.428 | 0.668 |
| | D:TreatmentAmbient + 2C:Sampling3 | -0.720 | 0.561 | -1.283 | 0.200 |
| | D:TreatmentAmbient + Periodic Heat Wave:Sampling3 | -0.181 | 0.561 | -0.322 | 0.747 |
| | D:TreatmentAmbient + 2C + Periodic Heat Wave:Sampling3 | -0.266 | 0.561 | -0.474 | 0.635 |
| **LAP** | (Intercept) | 5.799 | 0.212 | 27.380 | <0.001 |
| | D | 0.095 | 0.183 | 0.518 | 0.604 |
| | TreatmentAmbient + 2C | -0.347 | 0.183 | -1.900 | 0.057 |
| | TreatmentAmbient + Periodic Heat Wave | 0.158 | 0.183 | 0.864 | 0.388 |
| | TreatmentAmbient + 2C + Periodic Heat Wave | -0.207 | 0.183 | -1.135 | 0.256 |
| | Sampling2 | 0.276 | 0.183 | 1.513 | 0.130 |
| | Sampling3 | 0.008 | 0.183 | 0.046 | 0.964 |
| | D:TreatmentAmbient + 2C | 0.444 | 0.258 | 1.720 | 0.085 |
| | D:TreatmentAmbient + Periodic Heat Wave | -0.008 | 0.258 | -0.033 | 0.974 |
| | D:TreatmentAmbient + 2C + Periodic Heat Wave | 0.245 | 0.258 | 0.947 | 0.344 |
| | D:Sampling2 | 0.089 | 0.258 | 0.345 | 0.730 |
| | D:Sampling3 | -0.057 | 0.258 | -0.220 | 0.826 |
| | TreatmentAmbient + 2C:Sampling2 | 0.101 | 0.258 | 0.391 | 0.696 |
| | TreatmentAmbient + Periodic Heat Wave:Sampling2 | -0.232 | 0.258 | -0.897 | 0.369 |
| | TreatmentAmbient + 2C + Periodic Heat Wave:Sampling2 | 0.220 | 0.258 | 0.852 | 0.394 |
| | TreatmentAmbient + 2C:Sampling3 | 0.237 | 0.258 | 0.918 | 0.359 |
| | TreatmentAmbient + Periodic Heat Wave:Sampling3 | -0.131 | 0.258 | -0.507 | 0.612 |
| | TreatmentAmbient + 2C + Periodic Heat Wave:Sampling3 | 0.172 | 0.258 | 0.665 | 0.506 |
| | D:TreatmentAmbient + 2C:Sampling2 | -0.105 | 0.365 | -0.286 | 0.775 |
| | D:TreatmentAmbient + Periodic Heat Wave:Sampling2 | -0.046 | 0.365 | -0.126 | 0.900 |
| | D:TreatmentAmbient + 2C + Periodic Heat Wave:Sampling2 | -0.237 | 0.365 | -0.650 | 0.516 |
| | D:TreatmentAmbient + 2C:Sampling3 | -0.372 | 0.365 | -1.018 | 0.309 |
| | D:TreatmentAmbient + Periodic Heat Wave:Sampling3 | -0.165 | 0.365 | -0.452 | 0.651 |
| | D:TreatmentAmbient + 2C + Periodic Heat Wave:Sampling3 | -0.236 | 0.365 | -0.646 | 0.518 |
| **PHO** | (Intercept) | 6.178 | 0.180 | 34.313 | <0.001 |
| | D | 0.028 | 0.226 | 0.124 | 0.901 |
| | TreatmentAmbient + 2C | -0.305 | 0.231 | -1.318 | 0.187 |
| | TreatmentAmbient + Periodic Heat Wave | 0.140 | 0.231 | 0.606 | 0.545 |
| | TreatmentAmbient + 2C + Periodic Heat Wave | -0.158 | 0.226 | -0.701 | 0.483 |
| | Sampling2 | -0.117 | 0.222 | -0.528 | 0.597 |
| | Sampling3 | 0.090 | 0.222 | 0.406 | 0.685 |
| | D:TreatmentAmbient + 2C | 0.396 | 0.323 | 1.226 | 0.220 |
| | D:TreatmentAmbient + Periodic Heat Wave | -0.048 | 0.323 | -0.148 | 0.882 |

| | | | | |
|---|---|---|---|---|
| D:TreatmentAmbient + 2C + Periodic Heat Wave | -0.469 | 0.315 | -1.489 | 0.136 |
| D:Sampling2 | 0.187 | 0.309 | 0.605 | 0.545 |
| D:Sampling3 | -0.055 | 0.309 | -0.177 | 0.859 |
| TreatmentAmbient + 2C:Sampling2 | 0.093 | 0.313 | 0.297 | 0.767 |
| TreatmentAmbient + Periodic Heat Wave:Sampling2 | -0.259 | 0.313 | -0.826 | 0.409 |
| TreatmentAmbient + 2C + Periodic Heat Wave:Sampling2 | -0.024 | 0.309 | -0.078 | 0.938 |
| TreatmentAmbient + 2C:Sampling3 | 0.335 | 0.313 | 1.071 | 0.284 |
| TreatmentAmbient + Periodic Heat Wave:Sampling3 | -0.123 | 0.313 | -0.393 | 0.694 |
| TreatmentAmbient + 2C + Periodic Heat Wave:Sampling3 | 0.230 | 0.309 | 0.744 | 0.457 |
| D:TreatmentAmbient + 2C:Sampling2 | -0.150 | 0.440 | -0.340 | 0.734 |
| D:TreatmentAmbient + Periodic Heat Wave:Sampling2 | 0.055 | 0.440 | 0.126 | 0.900 |
| D:TreatmentAmbient + 2C + Periodic Heat Wave:Sampling2 | 0.294 | 0.434 | 0.678 | 0.498 |
| D:TreatmentAmbient + 2C:Sampling3 | -0.468 | 0.440 | -1.062 | 0.288 |
| D:TreatmentAmbient + Periodic Heat Wave:Sampling3 | 0.076 | 0.440 | 0.173 | 0.862 |
| D:TreatmentAmbient + 2C + Periodic Heat Wave:Sampling3 | 0.446 | 0.434 | 1.027 | 0.304 |

**Supplementary Table 4.** Output of the mixed-effect model to quantify the effects of drought on nutrient acquisition enzymatic ratios across warming regimes at 3 different sampling points. A model was run per each enzymatic ratio, it included an interaction effect of Drought (D), Treatment (warming regimes), and Sampling (Sampling 1 = Drought legacy, Sampling 2 = Resistance, Sampling 3 = Recovery). It counted for spatial and yearly variation by including Plot and Year as random effects. Enzymatic ratios are shown as follows: C:N = log (BG/(NAG+LAP)); N:P = log ((NAG+LAP)/PHO); C:P = log (BG/PHO); Ox:Hy =log ((POX + PER) / BG). Significant effects are marked in bold. SE stands for standard error.

| Nutrient acquisition enzymatic ratios | Predictor | Estimate | SE | Z-value | p-value |
|---|---|---|---|---|---|
| **C:N** | (Intercept) | -1.23059 | 0.163448 | -7.5289 | <0.001 |
| | D | -0.03429 | 0.215056 | -0.15944 | 0.8733 |
| | TreatmentAmbient + 2C | 0.222206 | 0.220607 | 1.007246 | 0.3138 |
| | TreatmentAmbient + Periodic Heat Wave | 0.426539 | 0.220607 | 1.933473 | 0.0532 |
| | TreatmentAmbient + 2C + Periodic Heat Wave | 0.314704 | 0.215056 | 1.46336 | 0.1434 |
| | **Sampling2** | **-0.56362** | **0.215056** | **-2.62079** | **0.0088** |
| | **Sampling3** | **0.453631** | **0.215056** | **2.109362** | **0.0349** |
| | D:TreatmentAmbient + 2C | -0.1423 | 0.308086 | -0.46187 | 0.6442 |
| | D:TreatmentAmbient + Periodic Heat Wave | -0.17869 | 0.308086 | -0.58 | 0.5619 |
| | D:TreatmentAmbient + 2C + Periodic Heat Wave | 0.078618 | 0.300083 | 0.261989 | 0.7933 |
| | D:Sampling2 | 0.503776 | 0.300083 | 1.678787 | 0.0932 |
| | D:Sampling3 | -0.01084 | 0.300083 | -0.03611 | 0.9712 |
| | TreatmentAmbient + 2C:Sampling2 | 0.135091 | 0.304086 | 0.444252 | 0.6569 |
| | TreatmentAmbient + Periodic Heat Wave:Sampling2 | -0.32592 | 0.304086 | -1.0718 | 0.2838 |
| | TreatmentAmbient + 2C + Periodic Heat Wave:Sampling2 | -0.04172 | 0.300083 | -0.13902 | 0.8894 |
| | TreatmentAmbient + 2C:Sampling3 | -0.23499 | 0.304086 | -0.77278 | 0.4397 |
| | TreatmentAmbient + Periodic Heat Wave:Sampling3 | -0.48404 | 0.304086 | -1.59177 | 0.1114 |
| | TreatmentAmbient + 2C + Periodic Heat Wave:Sampling3 | -0.07886 | 0.300083 | -0.26279 | 0.7927 |
| | D:TreatmentAmbient + 2C:Sampling2 | -0.23724 | 0.427222 | -0.55531 | 0.5787 |
| | D:TreatmentAmbient + Periodic Heat Wave:Sampling2 | 0.108791 | 0.427222 | 0.254647 | 0.7990 |
| | D:TreatmentAmbient + 2C + Periodic Heat Wave:Sampling2 | -0.51172 | 0.421488 | -1.21409 | 0.2247 |
| | D:TreatmentAmbient + 2C:Sampling3 | 0.03392 | 0.427222 | 0.079396 | 0.9367 |
| | D:TreatmentAmbient + Periodic Heat Wave:Sampling3 | 0.375737 | 0.427222 | 0.879489 | 0.3791 |
| | D:TreatmentAmbient + 2C + Periodic Heat Wave:Sampling3 | -0.17695 | 0.421488 | -0.41983 | 0.6746 |
| **C:P** | (Intercept) | -1.53161 | 0.210295 | -7.28315 | <0.001 |
| | D | -0.00593 | 0.178867 | -0.03317 | 0.9735 |
| | TreatmentAmbient + 2C | 0.159621 | 0.18349 | 0.869915 | 0.3843 |
| | TreatmentAmbient + Periodic Heat Wave | 0.355046 | 0.18349 | 1.934957 | 0.0530 |
| | TreatmentAmbient + 2C + Periodic Heat Wave | 0.23534 | 0.178867 | 1.315726 | 0.1883 |
| | Sampling2 | -0.22477 | 0.178867 | -1.2566 | 0.2089 |
| | Sampling3 | 0.342553 | 0.178867 | 1.915127 | 0.0555 |
| | D:TreatmentAmbient + 2C | -0.11617 | 0.256246 | -0.45335 | 0.6503 |
| | D:TreatmentAmbient + Periodic Heat Wave | -0.11363 | 0.256246 | -0.44343 | 0.6575 |
| | D:TreatmentAmbient + 2C + Periodic Heat Wave | 0.150439 | 0.252989 | 0.594648 | 0.5521 |
| | D:Sampling2 | 0.44687 | 0.24959 | 1.790414 | 0.0734 |
| | D:Sampling3 | 0.021627 | 0.24959 | 0.086649 | 0.9310 |
| | TreatmentAmbient + 2C:Sampling2 | 0.165571 | 0.252924 | 0.654628 | 0.5127 |
| | TreatmentAmbient + Periodic Heat Wave:Sampling2 | -0.20788 | 0.252924 | -0.8219 | 0.4111 |
| | TreatmentAmbient + 2C + Periodic Heat Wave:Sampling2 | 0.235837 | 0.24959 | 0.944898 | 0.3447 |

| | | | | | |
|---|---|---|---|---|---|
| | TreatmentAmbient + 2C:Sampling3 | -0.31803 | 0.252924 | -1.25742 | 0.2086 |
| | TreatmentAmbient + Periodic Heat Wave:Sampling3 | -0.40864 | 0.252924 | -1.61565 | 0.1062 |
| | TreatmentAmbient + 2C + Periodic Heat Wave:Sampling3 | -0.11218 | 0.24959 | -0.44947 | 0.6531 |
| | D:TreatmentAmbient + 2C:Sampling2 | -0.17309 | 0.355339 | -0.48713 | 0.6262 |
| | D:TreatmentAmbient + Periodic Heat Wave:Sampling2 | -0.02161 | 0.355339 | -0.06082 | 0.9515 |
| | D:TreatmentAmbient + 2C + Periodic Heat Wave:Sampling2 | -0.39982 | 0.352997 | -1.13263 | 0.2574 |
| | D:TreatmentAmbient + 2C:Sampling3 | 0.151097 | 0.355339 | 0.425218 | 0.6707 |
| | D:TreatmentAmbient + Periodic Heat Wave:Sampling3 | 0.116417 | 0.355339 | 0.327622 | 0.7432 |
| | D:TreatmentAmbient + 2C + Periodic Heat Wave:Sampling3 | -0.21417 | 0.352997 | -0.60673 | 0.5440 |
| **N:P** | (Intercept) | -0.2984 | 0.148349 | -2.01149 | 0.0443 |
| | D | 0.025735 | 0.140927 | 0.182613 | 0.8551 |
| | TreatmentAmbient + 2C | -0.06258 | 0.14457 | -0.4329 | 0.6651 |
| | TreatmentAmbient + Periodic Heat Wave | -0.07149 | 0.14457 | -0.49452 | 0.6209 |
| | TreatmentAmbient + 2C + Periodic Heat Wave | -0.08199 | 0.140927 | -0.58176 | 0.5607 |
| | **Sampling2** | **0.33623** | **0.140927** | **2.38584** | **0.0170** |
| | Sampling3 | -0.1137 | 0.140927 | -0.8068 | 0.4198 |
| | D:TreatmentAmbient + 2C | 0.02875 | 0.201893 | 0.142402 | 0.8868 |
| | D:TreatmentAmbient + Periodic Heat Wave | 0.067684 | 0.201893 | 0.335248 | 0.7374 |
| | D:TreatmentAmbient + 2C + Periodic Heat Wave | 0.09482 | 0.199327 | 0.475701 | 0.6343 |
| | D:Sampling2 | -0.05428 | 0.196649 | -0.27605 | 0.7825 |
| | D:Sampling3 | 0.035084 | 0.196649 | 0.17841 | 0.8584 |
| | TreatmentAmbient + 2C:Sampling2 | 0.03048 | 0.199275 | 0.152952 | 0.8784 |
| | TreatmentAmbient + Periodic Heat Wave:Sampling2 | 0.118041 | 0.199275 | 0.59235 | 0.5536 |
| | TreatmentAmbient + 2C + Periodic Heat Wave:Sampling2 | 0.280176 | 0.196649 | 1.424749 | 0.1542 |
| | TreatmentAmbient + 2C:Sampling3 | -0.08304 | 0.199275 | -0.41672 | 0.6769 |
| | TreatmentAmbient + Periodic Heat Wave:Sampling3 | 0.075399 | 0.199275 | 0.378365 | 0.7052 |
| | TreatmentAmbient + 2C + Periodic Heat Wave:Sampling3 | -0.0307 | 0.196649 | -0.15612 | 0.8759 |
| | D:TreatmentAmbient + 2C:Sampling2 | 0.061526 | 0.279967 | 0.219763 | 0.8261 |
| | D:TreatmentAmbient + Periodic Heat Wave:Sampling2 | -0.13302 | 0.279967 | -0.47514 | 0.6347 |
| | D:TreatmentAmbient + 2C + Periodic Heat Wave:Sampling2 | 0.088908 | 0.278122 | 0.319673 | 0.7492 |
| | D:TreatmentAmbient + 2C:Sampling3 | 0.114556 | 0.279967 | 0.409175 | 0.6824 |
| | D:TreatmentAmbient + Periodic Heat Wave:Sampling3 | -0.26194 | 0.279967 | -0.93561 | 0.3495 |
| | D:TreatmentAmbient + 2C + Periodic Heat Wave:Sampling3 | -0.06022 | 0.278122 | -0.21653 | 0.8286 |
| **Hy:Ox** | (Intercept) | 4.980747 | 0.138078 | 36.0721 | <0.001 |
| | D | -0.12903 | 0.166234 | -0.77619 | 0.4376 |
| | TreatmentAmbient + 2C | -0.19445 | 0.170481 | -1.14057 | 0.2540 |
| | TreatmentAmbient + Periodic Heat Wave | 0.19461 | 0.170482 | 1.141525 | 0.2537 |
| | TreatmentAmbient + 2C + Periodic Heat Wave | -0.10997 | 0.166234 | -0.66155 | 0.5083 |
| | Sampling2 | -0.19492 | 0.165962 | -1.17451 | 0.2402 |
| | Sampling3 | -0.07401 | 0.165962 | -0.44593 | 0.6556 |
| | D:TreatmentAmbient + 2C | 0.301307 | 0.238124 | 1.265334 | 0.2058 |
| | D:TreatmentAmbient + Periodic Heat Wave | 0.023866 | 0.238102 | 0.100235 | 0.9202 |
| | D:TreatmentAmbient + 2C + Periodic Heat Wave | 0.203338 | 0.23194 | 0.876682 | 0.3807 |
| | D:Sampling2 | 0.385216 | 0.23155 | 1.663642 | 0.0962 |
| | D:Sampling3 | 0.029974 | 0.23155 | 0.129449 | 0.8970 |
| | TreatmentAmbient + 2C:Sampling2 | 0.104052 | 0.234618 | 0.443494 | 0.6574 |

| | | | | |
|---|---|---|---|---|
| TreatmentAmbient + Periodic Heat Wave:Sampling2 | -0.23719 | 0.234618 | -1.01094 | 0.3120 |
| TreatmentAmbient + 2C + Periodic Heat Wave:Sampling2 | 0.079519 | 0.23155 | 0.343419 | 0.7313 |
| TreatmentAmbient + 2C:Sampling3 | 0.089996 | 0.234618 | 0.383587 | 0.7013 |
| TreatmentAmbient + Periodic Heat Wave:Sampling3 | -0.13609 | 0.234618 | -0.58006 | 0.5619 |
| TreatmentAmbient + 2C + Periodic Heat Wave:Sampling3 | 0.03416 | 0.23155 | 0.147526 | 0.8827 |
| D:TreatmentAmbient + 2C:Sampling2 | -0.12788 | 0.329646 | -0.38793 | 0.6981 |
| D:TreatmentAmbient + Periodic Heat Wave:Sampling2 | -0.11667 | 0.329629 | -0.35393 | 0.7234 |
| D:TreatmentAmbient + 2C + Periodic Heat Wave:Sampling2 | -0.37917 | 0.325207 | -1.16594 | 0.2436 |
| D:TreatmentAmbient + 2C:Sampling3 | -0.11879 | 0.329646 | -0.36037 | 0.7186 |
| D:TreatmentAmbient + Periodic Heat Wave:Sampling3 | -0.09434 | 0.329629 | -0.28619 | 0.7747 |
| D:TreatmentAmbient + 2C + Periodic Heat Wave:Sampling3 | -0.03179 | 0.325207 | -0.09777 | 0.9221 |

**Supplementary Table 5.** Output of the mixed-effect model to quantify the effects of drought on Glycogen production, and Dehydrogenase activity warming regimes at 3 different sampling points. A model was run for Glycogen and a second one for Dehydrogenase, and each model included an interaction effect of Drought (D), Treatment (warming regimes), and Sampling point (Sampling 1 = Drought legacy, Sampling 2 = Resistance, Sampling 3 = Recovery). It counted for spatial variation as random effect. Significant effects are marked in bold. Gly = Glycogen, DEH = dehydrogenase. SE stands for standard error.

| Class | Predictor | Estimate | SE | Z-value | p-value |
|---|---|---|---|---|---|
| **GLY** | (Intercept) | 0.074 | 0.021742 | 3.403483 | <0.001 |
| | Drought | 0.104 | 0.030748 | 3.382286 | **0.000719** |
| | TreatmentAmbient + 2C | 0.046 | 0.030748 | 1.496012 | 0.134651 |
| | TreatmentAmbient + Periodic Heat Wave | 2.75E-08 | 0.030748 | 8.95E-07 | 0.999999 |
| | TreatmentAmbient + 2C + Periodic Heat Wave | -0.004 | 0.030748 | -0.13009 | 0.896498 |
| | Sampling2 | -0.008 | 0.030748 | -0.26018 | 0.794729 |
| | Sampling3 | 0.216 | 0.030748 | 7.024747 | <0.001 |
| | Drought:TreatmentAmbient + 2C | -0.05 | 0.043485 | -1.14983 | 0.250216 |
| | Drought:TreatmentAmbient + Periodic Heat Wave | -0.098 | 0.043485 | -2.25366 | 0.024218 |
| | Drought:TreatmentAmbient + 2C + Periodic Heat Wave | -0.082 | 0.043485 | -1.88571 | 0.059333 |
| | Drought:Sampling2 | -0.12 | 0.043485 | -2.75958 | 0.005788 |
| | Drought:Sampling3 | -0.078 | 0.043485 | -1.79373 | 0.072857 |
| | TreatmentAmbient + 2C:Sampling2 | -0.04 | 0.043485 | -0.91986 | 0.357645 |
| | TreatmentAmbient + Periodic Heat Wave:Sampling2 | 0.002 | 0.043485 | 0.045992 | 0.963317 |
| | TreatmentAmbient + 2C + Periodic Heat Wave:Sampling2 | -0.014 | 0.043485 | -0.32195 | 0.747489 |
| | TreatmentAmbient + 2C:Sampling3 | -0.098 | 0.043485 | -2.25366 | **0.024218** |
| | TreatmentAmbient + Periodic Heat Wave:Sampling3 | -0.036 | 0.043485 | -0.82788 | 0.407741 |
| | TreatmentAmbient + 2C + Periodic Heat Wave:Sampling3 | -0.032 | 0.043485 | -0.73589 | 0.461798 |
| | Drought:TreatmentAmbient + 2C:Sampling2 | 0.056 | 0.061497 | 0.910615 | 0.362498 |
| | Drought:TreatmentAmbient + Periodic Heat Wave:Sampling2 | 0.088 | 0.061497 | 1.430968 | 0.152439 |
| | Drought:TreatmentAmbient + 2C + Periodic Heat Wave:Sampling2 | 0.08 | 0.061497 | 1.300879 | 0.1933 |
| | Drought:TreatmentAmbient + 2C:Sampling3 | 0.06 | 0.061497 | 0.975659 | 0.329233 |
| | Drought:TreatmentAmbient + Periodic Heat Wave:Sampling3 | 0.092 | 0.061497 | 1.496012 | 0.134651 |
| | DroughtC:TreatmentAmbient + 2C + Periodic Heat Wave:Sampling3 | 0.088 | 0.061497 | 1.430967 | 0.15244 |
| **DEH** | (Intercept) | 0.118 | 0.00964 | 12.24042 | <0.001 |
| | Drought | 0.04 | 0.013633 | 2.934002 | **0.003346** |
| | TreatmentAmbient + 2C | 0.014 | 0.013633 | 1.026883 | 0.304476 |
| | TreatmentAmbient + Periodic Heat Wave | 0.002 | 0.013633 | 0.146694 | 0.883373 |
| | TreatmentAmbient + 2C + Periodic Heat Wave | 0.006 | 0.013633 | 0.440114 | 0.659855 |
| | Sampling2 | -0.034 | 0.011524 | -2.95039 | **0.003174** |
| | Sampling3 | -0.044 | 0.011524 | -3.81815 | **0.000134** |
| | Drought:TreatmentAmbient + 2C | -0.038 | 0.01928 | -1.97091 | **0.048734** |
| | Drought:TreatmentAmbient + Periodic Heat Wave | -0.042 | 0.01928 | -2.17837 | **0.029379** |
| | Drought:TreatmentAmbient + 2C + Periodic Heat Wave | -0.038 | 0.01928 | -1.97093 | **0.048732** |
| | Drought:Sampling2 | -0.022 | 0.016297 | -1.34993 | 0.177039 |
| | Drought:Sampling3 | -0.036 | 0.016297 | -2.20898 | **0.027176** |
| | TreatmentAmbient + 2C:Sampling2 | -0.014 | 0.016297 | -0.85904 | 0.39032 |
| | TreatmentAmbient + Periodic Heat Wave:Sampling2 | -0.012 | 0.016297 | -0.7363 | 0.461547 |
| | TreatmentAmbient + 2C + Periodic Heat Wave:Sampling2 | -0.022 | 0.016297 | -1.34992 | 0.177042 |
| | TreatmentAmbient + 2C:Sampling3 | -0.03 | 0.016297 | -1.8408 | 0.065651 |
| | TreatmentAmbient + Periodic Heat Wave:Sampling3 | -0.01 | 0.016297 | -0.6136 | 0.539479 |
| | TreatmentAmbient + 2C + Periodic Heat Wave:Sampling3 | -0.01 | 0.016297 | -0.6136 | 0.539479 |

| | | | | |
|---|---|---|---|---|
| Drought:TreatmentAmbient + 2C:Sampling2 | 0.028 | 0.023048 | 1.214871 | 0.224415 |
| Drought:TreatmentAmbient + Periodic Heat Wave:Sampling2 | 0.037999 | 0.023048 | 1.648727 | 0.099204 |
| Drought:TreatmentAmbient + 2C + Periodic Heat Wave:Sampling2 | 0.038 | 0.023048 | 1.64873 | 0.099203 |
| Drought:TreatmentAmbient + 2C:Sampling3 | 0.05 | 0.023048 | 2.169406 | **0.030052** |
| Drought:TreatmentAmbient + Periodic Heat Wave:Sampling3 | 0.044 | 0.023048 | 1.909065 | 0.056254 |
| Drought:TreatmentAmbient + 2C + Periodic Heat Wave:Sampling3 | 0.046 | 0.023048 | 1.995845 | **0.045951** |

**Supplementary Table 6. Effects of extreme climate treatments, sampling phase, and microbial C-strategies on log response ratios (LRRs).** *C-process = extracellular hydrolytic (BG) and oxidative (POX + PER) C-acquisition, and internal C storage (GLY). Sampling = Drought legacy, Resistance, and Recovery. Extremes: Drought (D), Constant Warming (W), Heat Waves (HW), W + HW, D x W, D x HW, D x W + HW. Type II Wald $\chi^2$ tests from a generalised linear mixed model (gaussian family). Significance: *** $p < 0.001$, ** $p < 0.01$, * $p < 0.05$, . $p < 0.1$, ns = not significant. Random structure: unstructured covariance among trait types within plot.*

| **Factors** | $\chi^2$ | ***df*** | ***p*** | **significance** |
|---|---:|---:|---|---|
| **C-process** | **56.05** | **2** | **<0.001** | *** |
| Extreme | 6.91 | 6 | 0.329 | ns |
| Sampling | 5.22 | 2 | 0.073 | . |
| **C-process × Extreme** | **32.70** | **12** | **0.001** | ** |
| **C-process × Sampling** | **84.13** | **4** | **<0.001** | *** |
| Extreme × Sampling | 8.52 | 12 | 0.743 | ns |
| **C-process × Extreme × Sampling** | **55.00** | **24** | **<0.001** | *** |

**Supplementary Table 7.** Estimated marginal means of log response ratios (LRR) for microbial C acquisition and storage processes across climate extreme treatments and sampling phases. Estimated marginal means ± SE. 95% confidence intervals adjusted using the Bonferroni method. P-values adjusted using false discovery rate (FDR). 'vs. zero': − = significantly suppressed relative to control (95% CI below zero); + = significantly stimulated (95% CI above zero); ns = not significant. Groups sharing the same letter do not differ significantly (α = 0.05). BG = hydrolytic C-acquisition; POXPER = oxidative C-acquisition; GLY = C-storage.

| Phase | Treatment | Process | Mean LRR | SE | Lower 95% CI | Upper 95% CI | vs. zero | Group |
|---|---|---|---|---|---|---|---|---|
| Drought legacy | D | BG | 0.136 | 0.112 | -0.133 | 0.406 | ns | b |
| | | POXPER | -0.012 | 0.068 | -0.176 | 0.153 | ns | b |
| | | **GLY** | **-1.020** | **0.222** | **-1.555** | **-0.485** | **-** | **a** |
| | W | BG | -0.176 | 0.112 | -0.446 | 0.093 | ns | a |
| | | POXPER | 0.033 | 0.068 | -0.131 | 0.197 | ns | a |
| | | GLY | -0.160 | 0.222 | -0.695 | 0.375 | ns | a |
| | HW | BG | 0.117 | 0.112 | -0.153 | 0.386 | ns | b |
| | | POXPER | -0.037 | 0.068 | -0.202 | 0.127 | ns | b |
| | | **GLY** | **-1.236** | **0.222** | **-1.771** | **-0.701** | **-** | **a** |
| | W+HW | BG | 0.139 | 0.112 | -0.131 | 0.409 | ns | b |
| | | POXPER | -0.015 | 0.068 | -0.179 | 0.150 | ns | b |
| | | **GLY** | **-0.883** | **0.222** | **-1.418** | **-0.348** | **-** | **a** |
| | D*W | BG | -0.052 | 0.112 | -0.322 | 0.218 | ns | a |
| | | POXPER | -0.131 | 0.068 | -0.295 | 0.034 | ns | a |
| | | GLY | -0.478 | 0.222 | -1.013 | 0.057 | ns | a |
| | **D*HW** | **BG** | **0.258** | **0.112** | **-0.012** | **0.528** | **ns** | **c** |
| | | **POXPER** | **-0.098** | **0.068** | **-0.263** | **0.066** | **ns** | **b** |
| | | **GLY** | **-1.060** | **0.222** | **-1.595** | **-0.525** | **-** | **a** |
| | D*W+HW | BG | -0.074 | 0.112 | -0.344 | 0.196 | ns | b |
| | | POXPER | -0.159 | 0.068 | -0.324 | 0.005 | ns | b |
| | | **GLY** | **-1.084** | **0.222** | **-1.619** | **-0.549** | **-** | **a** |
| Resistance | D | **BG** | **-0.411** | **0.112** | **-0.681** | **-0.141** | **-** | **a** |
| | | **POXPER** | **0.197** | **0.068** | **0.032** | **0.361** | **+** | **b** |
| | | GLY | 0.005 | 0.222 | -0.530 | 0.540 | ns | ab |
| | W | BG | 0.006 | 0.112 | -0.264 | 0.276 | ns | a |
| | | POXPER | 0.055 | 0.068 | -0.109 | 0.220 | ns | a |
| | | GLY | 0.140 | 0.222 | -0.395 | 0.675 | ns | a |
| | HW | BG | 0.045 | 0.112 | -0.225 | 0.315 | ns | a |
| | | POXPER | 0.123 | 0.068 | -0.041 | 0.287 | ns | a |
| | | GLY | -0.249 | 0.222 | -0.784 | 0.286 | ns | a |
| | W+HW | BG | 0.085 | 0.112 | -0.184 | 0.355 | ns | b |
| | | POXPER | -0.012 | 0.068 | -0.176 | 0.153 | ns | b |
| | | **GLY** | **-0.673** | **0.222** | **-1.208** | **-0.138** | **-** | **a** |
| | D*W | **BG** | **-0.443** | **0.112** | **-0.713** | **-0.173** | **-** | **a** |
| | | POXPER | 0.008 | 0.068 | -0.156 | 0.172 | ns | b |
| | | GLY | 0.306 | 0.222 | -0.229 | 0.841 | ns | b |
| | D*HW | **BG** | **-0.440** | **0.112** | **-0.710** | **-0.170** | **-** | **a** |
| | | POXPER | 0.110 | 0.068 | -0.054 | 0.274 | ns | b |
| | | GLY | 0.258 | 0.222 | -0.277 | 0.793 | ns | b |
| | D*W+HW | BG | -0.247 | 0.112 | -0.517 | 0.023 | ns | a |
| | | POXPER | 0.011 | 0.068 | -0.153 | 0.175 | ns | a |
| | | GLY | -0.191 | 0.222 | -0.726 | 0.344 | ns | a |

| Phase | Treatment | Process | Mean LRR | SE | Lower 95% CI | Upper 95% CI | vs. zero | Group |
|---|---|---|---|---|---|---|---|---|
| Recovery | D | BG | 0.030 | 0.112 | -0.240 | 0.300 | ns | a |
| | | POXPER | -0.019 | 0.068 | -0.184 | 0.145 | ns | a |
| | | GLY | -0.103 | 0.222 | -0.638 | 0.432 | ns | a |
| | W | BG | -0.224 | 0.112 | -0.494 | 0.046 | ns | a |
| | | POXPER | -0.064 | 0.068 | -0.228 | 0.100 | ns | a |
| | | GLY | -0.169 | 0.222 | -0.704 | 0.366 | ns | a |
| | HW | BG | 0.041 | 0.112 | -0.229 | 0.311 | ns | a |
| | | POXPER | -0.032 | 0.068 | -0.196 | 0.132 | ns | a |
| | | GLY | -0.163 | 0.222 | -0.698 | 0.372 | ns | a |
| | W+HW | BG | -0.062 | 0.112 | -0.332 | 0.208 | ns | a |
| | | POXPER | -0.073 | 0.068 | -0.237 | 0.091 | ns | a |
| | | GLY | -0.125 | 0.222 | -0.660 | 0.410 | ns | a |
| | D*W | BG | -0.292 | 0.112 | -0.562 | -0.022 | - | a |
| | | POXPER | -0.001 | 0.068 | -0.166 | 0.163 | ns | a |
| | | GLY | -0.300 | 0.222 | -0.835 | 0.235 | ns | a |
| | D*HW | BG | -0.137 | 0.112 | -0.407 | 0.133 | ns | a |
| | | POXPER | -0.060 | 0.068 | -0.224 | 0.104 | ns | a |
| | | GLY | -0.235 | 0.222 | -0.770 | 0.300 | ns | a |
| | D*W+HW | BG | 0.058 | 0.112 | -0.212 | 0.328 | ns | a |
| | | POXPER | 0.020 | 0.068 | -0.144 | 0.185 | ns | a |
| | | GLY | -0.247 | 0.222 | -0.782 | 0.288 | ns | a |

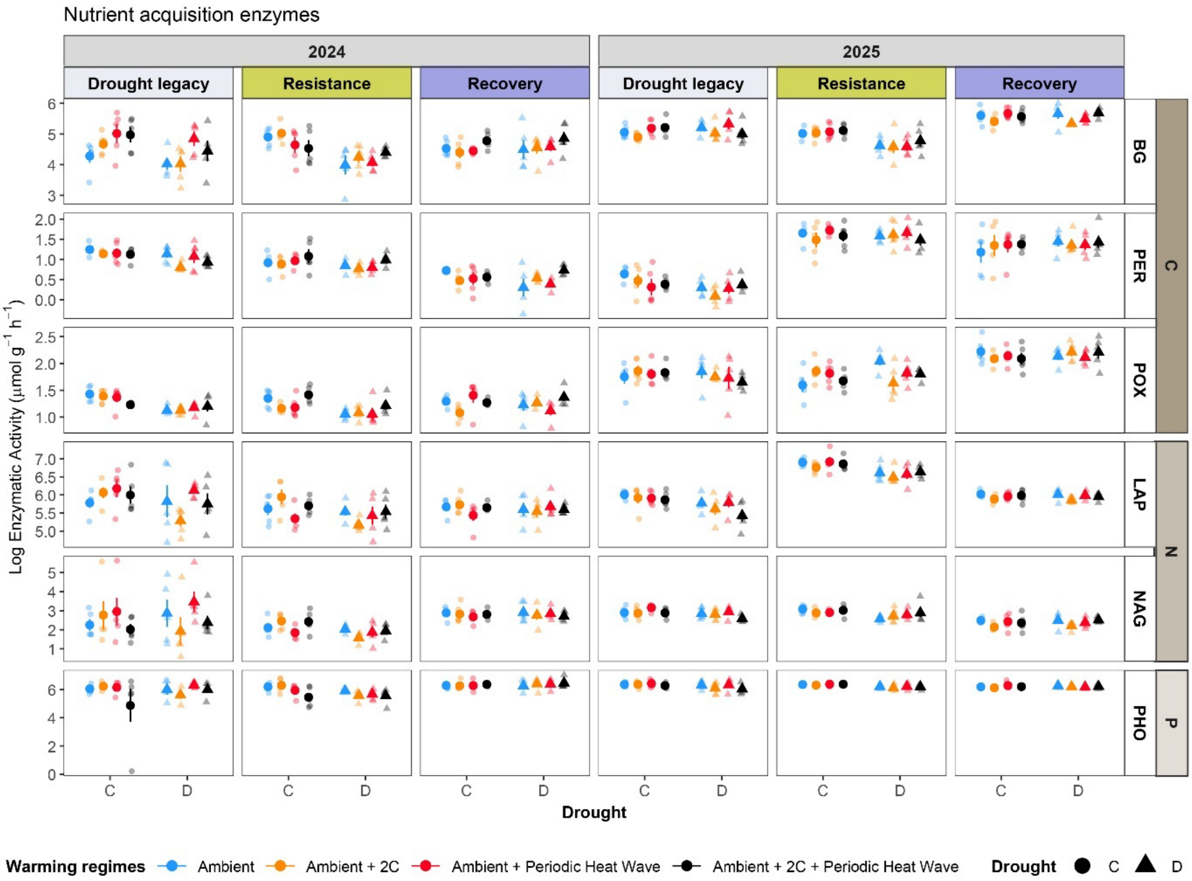


**Supplementary Figure 1.** Raw data of soil extracellular enzymes per year: 2024 and 2025. Samples were taken 3 times per year: Drought legacy phase, resistance phase, and recovery phase. Colours indicate warming regimes, blue shows drought effect with no warming, orange = drought effect modulated by 'W', red = drought effect modulated 'HW', black = drought effect modulated by 'W + HW'. Drought levels by shape: C = Control (circle); D = Drought (triangle).

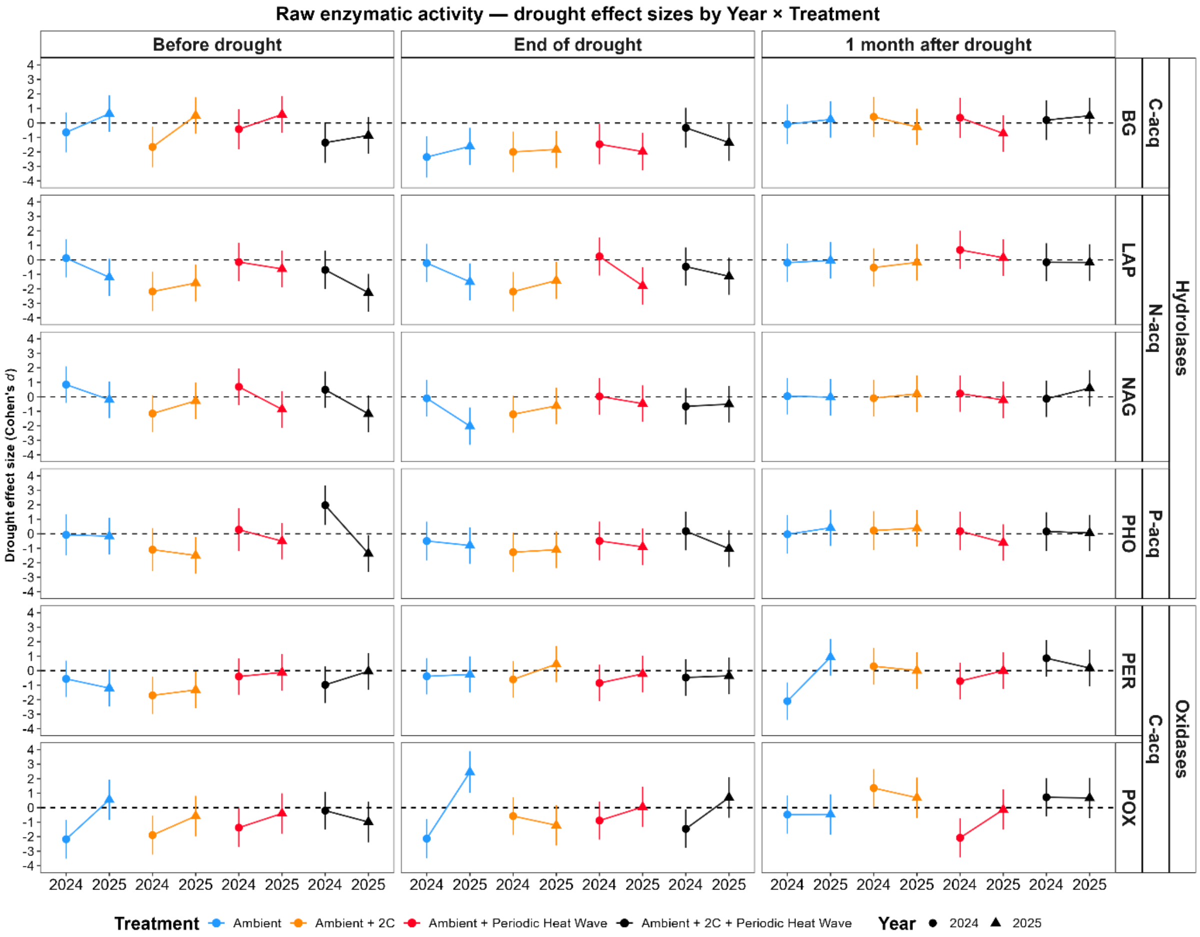


**Supplementary Figure 2.** Effect of drought on microbial nutrient acquisition across warming regimes. Effect sizes (Cohen's d ± 95% CI) represent the standardized difference in enzyme activity between drought and control plots under four warming treatments, measured at three time points for each year: 2024 and 2025: *Drought legacy* (before drought), *Resistance* (at the end of the drought), and *Recovery* (one month after rewetting) (n = 240). Panel are displayed by nutrient-association groups (C-, N-, or P-acquiring enzymes). Individual enzymes are as follows: PER = peroxidase; POX = phenol oxidase; BG = β-glucosidase; LAP = leucine aminopeptidase; NAG = β-N-acetylglucosaminidase; PHO = phosphatase. Colours indicate warming regimes, blue shows drought effect with no warming, orange = drought effect modulated by 'W', red = drought effect modulated 'HW', black = drought effect modulated by 'W + HW'.

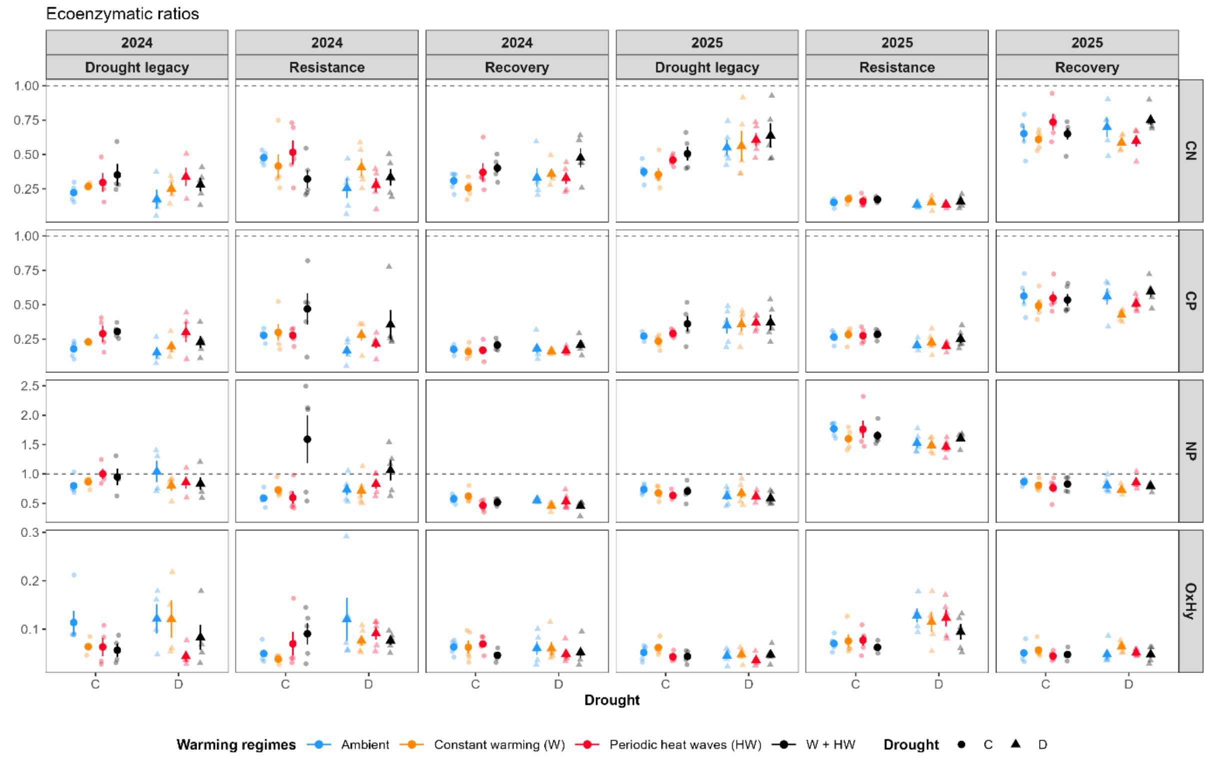


**Supplementary Figure 3.** Raw data of ecoenzymatic stoichiometry at each year: 2024 and 2025. Colours indicate warming regimes, blue shows drought effect with no warming, orange = drought effect modulated by 'W', red = drought effect modulated 'HW', black = drought effect modulated by 'W + HW'. Drought levels by shape: C = Control (circle); D = Drought (triangle).

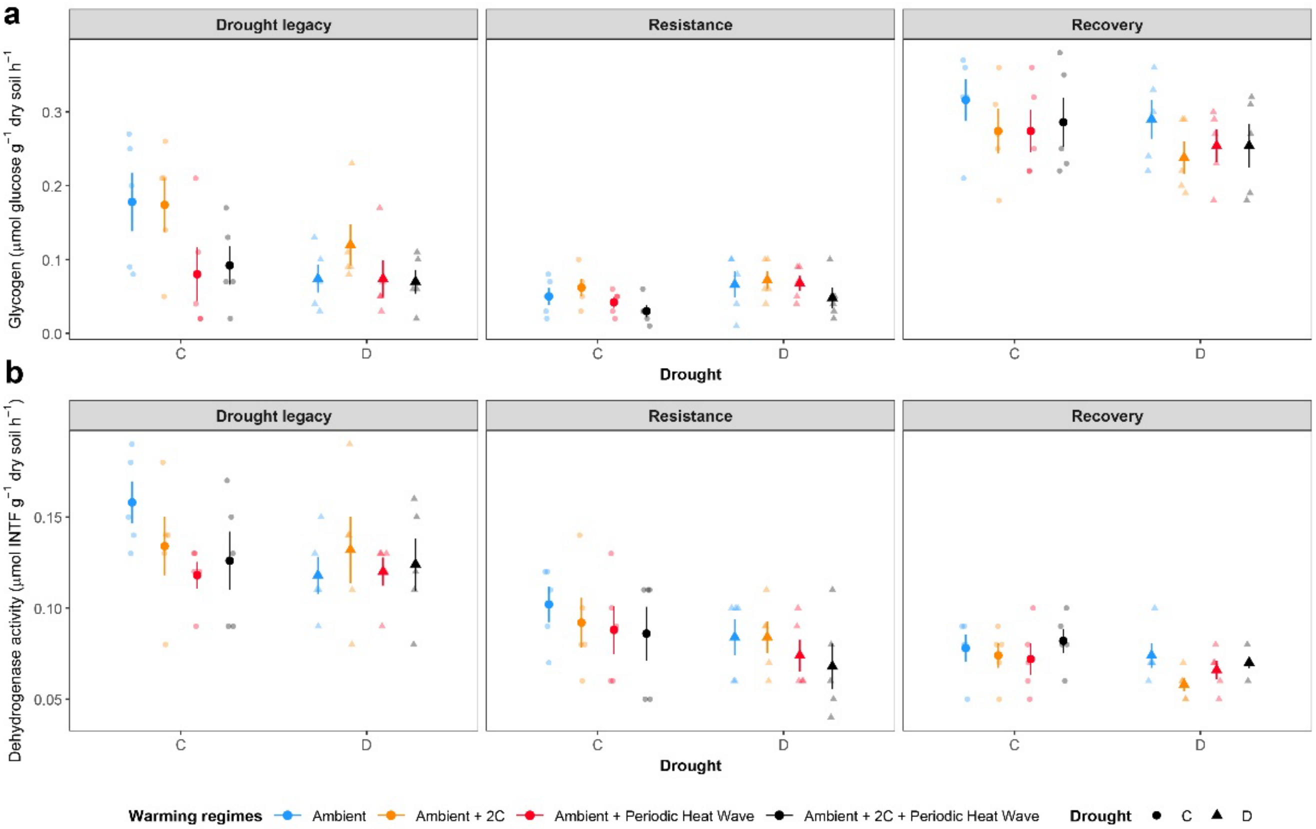


**Supplementary Figure 4.** Raw data of glycogen concentration and dehydrogenase activity at each sampling phase: Drought legacy, resistance, and recovery. Colours indicate warming regimes, blue shows drought effect with no warming, orange = drought effect modulated by 'W', red = drought effect modulated 'HW', black = drought effect modulated by 'W + HW'. Drought levels by shape: C = Control (circle); D = Drought (triangle).
